# Ibrutinib blocks YAP1 activation and reverses BRAFi resistance in melanoma cells

**DOI:** 10.1101/2020.03.25.006916

**Authors:** SA Misek, PA Newbury, E Chekalin, S Paithankar, AI Doseff, B Chen, KA Gallo, RR Neubig

## Abstract

Most BRAF-mutant melanoma tumors respond initially to BRAFi/MEKi therapy, although few patients have durable long-term responses to these agents. The goal of this study was to utilize an unbiased computational approach to identify inhibitors which reverse an experimentally derived BRAFi resistance gene expression signature. Using this approach, we found that ibrutinib effectively reverses this signature and we demonstrate experimentally that ibrutinib re-sensitizes a subset of BRAFi-resistant melanoma cells to vemurafenib. Ibrutinib is used clinically as a BTK inhibitor; however, neither BTK deletion nor treatment with acalabrutinib, another BTK inhibitor with reduced off-target activity, re-sensitized cells to vemurafenib. These data suggest that ibrutinib acts through a BTK-independent mechanism in vemurafenib re-sensitization. To better understand this mechanism, we analyzed the transcriptional profile of ibrutinib-treated BRAFi-resistant melanoma cells and found that the transcriptional profile of ibrutinib was highly similar to that of multiple SRC kinase inhibitors. Since ibrutinib, but not acalabrutinib, has significant off-target activity against multiple SRC family kinases, it suggests that ibrutinib may be acting through this mechanism. Furthermore, genes either upregulated or downregulated by ibrutinib treatment are enriched in YAP1 target genes and we showed that ibrutinib, but not acalabrutinib, reduces YAP1 activity in BRAFi-resistant melanoma cells. Taken together, these data suggest that ibrutinib, or other SRC family kinase inhibitors, may be useful for treating some BRAFi/MEKi-refractory melanoma tumors.

## Introduction

Approximately 90% of melanoma tumors harbor activating mutations in the MAPK pathway and most of these tumors have BRAF^V600^ mutations (1). Most BRAF-mutant melanoma tumors initially respond to BRAF inhibitors (BRAFi), however, this response is often short-lived and most tumors develop resistance (2,3). Mechanisms of resistance to BRAFi/MEKi therapy most commonly occur through re-activation of the mitogen activated protein kinase (MAPK) pathway (4–20). However, there are few if any effective clinical interventions that overcome BRAFi resistance after it develops. In this study, we sought to identify compounds which reverse a BRAFi resistance gene signature. This systems-based approach has been widely explored in cancer drug discovery (21–25), yet few studies have investigated resistance in melanoma. Ultimately, the goal is to identify drugs which could be combined with BRAFi/MEKi therapy to prevent or reverse drug resistance.

One advantage to using this approach is that it allows for the identification of compounds whose effects may result from complex polypharmacology. There are several examples of the clinical utility of drugs that exhibit polypharmacology, including crizotinib, afatinib, ceritinib, dasatinib, erlotinib, nilotinib, ponatinib, and imatinib (26). In the case of imatinib, it was first developed to inhibit a BCR-ABL fusion protein in CML (27–29). But later imatinib was used to target dermatofibrosarcoma protuberans tumors harboring gene fusions which result in aberrant PDGFR activation or gastrointestinal stromal tumors which have activating PDGFRA or KIT mutations since imatinib has off-target activity against PDGFR and KIT (30–35). Several molecules, many of which are not kinase inhibitors, are currently under clinical investigation and have a mechanism of action linked to previously unappreciated off-target effects (36). These examples likely represent only a fraction of circumstances in which kinase inhibitor polypharmacology is clinically relevant. Because of this, there have been recent large-scale efforts to profile kinase inhibitor polypharmacology (37). Defining the entire polypharmacology network will result in a sizeable increase in the number of clinically actionable applications.

In this study we identify a new role for ibrutinib, an FDA-approved BTK inhibitor, in reversing BRAFi resistance in melanoma *in silico* and *in vitro*. Our studies suggest that ibrutinib may modulate YAP1 activation in BRAFi resistant melanoma cells. YAP1 is a transcriptional co-activator whose activity is regulated by the actin cytoskeleton, as well as through changes in the phosphorylation state of YAP1 (38–41). Some phosphorylation events on YAP1 by LATS1/2 lead to inactivation and subsequent proteasomal degradation (42) whereas phosphorylation at other sites, targeted by YES1 and other kinases, is critical for YAP1 nuclear translocation and activation (43). YAP1 is activated in BRAFi-resistant melanoma cells and silencing or deletion of YAP1 reverses BRAFi resistance (4,44–47). In addition to melanoma, YAP1 has been implicated in many other cancer types including breast cancer (48), glioblastoma (49), pancreatic cancer (50), hepatocellular carcinoma (51), and non-small-cell lung cancer (52). Despite the importance of YAP1 in cancer, it is still difficult to pharmacologically target YAP1. Verteporfin, a drug used to treat macular degeneration, blocks YAP1-TEAD activity *in vitro*, but has limited efficacy *in vivo* (53). Since YAP1 activity is regulated by its phosphorylation state, it may be possible to utilize the polypharmacology of FDA-approved kinase inhibitors to indirectly block YAP1 activation. In this study we found that ibrutinib blocks the nuclear accumulation of YAP1, suggesting that it may be possible to re-purpose ibrutinib or related SRC-family kinase inhibitors to treat YAP1-driven cancers.

## Materials and Methods

### Cell lines, reagents, and antibodies

Parental (denoted by a *P* suffix in the cell line name) and matched isogenic BRAFi-resistant cells (denoted by an *R* suffix in the cell line name) were either a gift (M229P/R, M238P/R) from Dr. Roger Lo (UCLA) or generated in our laboratory (UACC62P/R). These cells were generated and cultured as described below (47).

Luteolin (#10004161), BVT-948 (#16615), ketoprofen (#10006661), lestaurtinib (#12094), L-NMMA (#10005031), ibrutinib (#16274), acalabrutinib (#19899), fadrozole (#24272), letrozole (#11568), exemestane (#15008), and vemurafenib (#10618) were purchased from Cayman Chemical (Ann Arbor, USA). Pyrvinium pamoate (#HY-A0293) was purchased from MedChemExpress (Monmouth Junction, USA). Clofilium tosylate (#C2365) was purchased from Sigma Aldrich (St. Louis, USA). All compounds (except L-NMMA) were diluted in DMSO to a stock concentration of 10 mM. L-NMMA was diluted in H_2_O to a stock concentration of 0.5 mM. All compounds were aliquoted and stored at −20°C.

Antibodies against YAP1 (#14074) and TAZ (#83669) were purchased from Cell Signaling (Danvers, USA). An antibody against Actin (#sc1616) was purchased from Santa Cruz Biotechnology (Dallas, USA). Donkey anti-Mouse800 (#926-32212), Donkey anti-Goat680 (#926-68074), and Donkey anti-Rabbit680 (#926-68073) immunoblotting secondary antibodies were purchased from LI-COR (Lincoln, USA). Anti-rabbit-HRP (#7074) immunoblotting secondary was purchased from Cell Signaling Technology. Alexa Fluor goat anti-rabbit488 (#A11034) and donkey anti-goat488 (A11055) were purchased from Invitrogen (Carlsbad, USA).

### Cell culture

Cells were cultured in DMEM (ThermoFisher, Waltham, USA #11995-065) supplemented with 10% FBS (ThermoFisher, #10437-028) and 1% Antibiotic-Antimycotic (ThermoFisher, #15240062) and were passaged at approximately 75% confluence. The BRAFi-resistant cell line variants were maintained in culture medium supplemented with 2 μM vemurafenib. Vemurafenib was removed from the culture medium when cells were seeded for experiments, except where otherwise indicated. Cells were routinely tested for mycoplasma contamination by DAPI staining. Short Tandem Repeat (STR) profiling on all cell lines was performed at the MSU genomics core. In all cases, isogenic pairs of cell lines had identical STR profiles. After thawing cells were used for either 2 months or 20 passages, whichever came first.

### Cloning/CRISPR

For CRISPR experiments the sgRNA were: sgControl (5’-TCCCCGAGACCATCTTAGGG-3’), sgBTK #1 (5’-ATGAGTATGACTTTGAACGT-3’), and sgBTK #2 (5’-CCCTTCATCATATACAACCT-3’). These guide sequences were cloned into pLentiCRISPRv2-Puro (from Feng Zhang, Addgene plasmid #52961). Successful cloning was confirmed by Sanger sequencing. To measure knockout efficiency, amplicons containing the CRISPR cut sites were amplified from the genomic DNA with PCR and the ratio of frameshifted/functional DNA species was measured with Sanger sequencing using the TIDE algorithm (54). The primers for gDNA amplification and Sanger sequencing are listed in (Table S1).

### Virus preparation and infection

HEK-293T cells were seeded into 10-cm plates at a density of 4×10^6^ cells/plate and the cells were allowed to attach overnight. The next day the cells were transfected with a plasmid cocktail containing 5000 ng of the pLentiCRISPRv2 plasmid, 5000 ng of psPAX2 (Addgene plasmid #12260), 500 ng of pMD2.G (Addgene plasmid #12259), and 20 μL of Lipofectamine 2000 (ThermoFisher, #11668019) in 400 μL of OptiMEM (ThermoFisher, #31985070). The next morning the medium was changed to 10 mL of fresh complete culture medium, and the following day each plate was supplemented with an additional 5 mL of culture medium. After 24 h, the culture medium was harvested and filtered through a 0.45-μm syringe filter. Virus was stored at 4°C and was used within 2 weeks.

Melanoma cells were seeded into 10-cm plates at a density of 5×10^5^ cells/plate in 10 mL of complete culture medium. While the cells were still in suspension, 3 mL of viral supernatant was added to each plate. The cells were incubated with virus overnight, then the medium was changed to 10 mL of fresh medium. After 24 h, the medium was changed to 10 mL of fresh medium supplemented with puromycin (2 μM). The cells were cultured in the presence of selection antibiotic until all the cells on the kill control plate died (approximately 3 days). Individual clones for the CRISPR cell lines were not selected, but instead we used a pooled infection approach. Validation of CRISPR knockout efficiency was performed by Sanger sequencing as described above.

### Viability experiments

Cells were seeded into 384-well tissue culture plates (PerkinElmer, Waltham, USA, #6007689) at a density of 1000 cells/well in 20 μL of growth medium. The next day, compounds were pre-diluted in growth medium then added to the 384-well plates so that the final volume of each well was 40 μL. A PBS or growth medium barrier was added to the outer wells of the plate to limit evaporation. Cells were cultured under these conditions for 72 h. To assess viability, 8 μL of CellTiter-Glo (Promega, Madison, USA, #G7573) was added to each well. Plates were incubated on orbital shaker for 5 min at room temperature, then briefly centrifuged (4000 rpm, 60 s) before being read on a Bio-Tek Synergy Neo plate reader with the #11 and #41 Ex/Em filter cubes. Viability signal is plotted versus log (Vemurafenib concentration) for each treatment condition.

### Flow cytometry

#### Cell cycle

Cells were rinsed once in PBS before being trypsinized, washed once in PBS and immediately fixed in 70% ethanol for 20 min at room temperature. The cells were washed once and were re-suspended in PBS supplemented with 20μg/mL propidium iodide (#P1304MP, ThermoFisher) and 200 μg/mL RNaseA. The cells were briefly mixed and were incubated on ice for 20 min. Following incubation, the cells were filtered through a 70 μM filter and were run on an Accuri C6 flow cytometer (BD Biosciences, Franklin Lakes, USA). Data were analyzed with the FCS Express flow cytometry analysis software package.

#### Annexin V/Propidium Iodide

Both floating and adherent cells were collected by trypsinization. The cells were pelleted, washed once in PBS, and then re-suspended in 200 μL of Annexin V binding buffer (10 mM HEPES pH 7.4, 140 mM NaCl, 2.5 mM CaCl_2_) and 1 μL of APC-conjugated Annexin V (ThermoFisher, #A35110) on ice in the dark for 20 min. The cells were pelleted and re-suspended in 500 μL Annexin V binding buffer with 2 μg/mL propidium iodide. After 20 min the cells were filtered through a 70 μM filter and were run on an Accuri C6 flow cytometer. Data were analyzed with the FCS Express flow cytometry analysis software package.

### DEVD Assay

Both the floating and attached cells were collected, rinsed as described above and then lysed in 200 μL of Triton-X100 lysis buffer (25 mM HEPES, 100 mM NaCl, 1 mM EDTA, 10% glycerol, 1% Triton X-100) supplemented with protease/phosphatase inhibitors. The lysates were centrifuged at 20,000g for 15 min. In a 384-well plate 10 μL of 2x Cytobuffer (100 mM PIPES pH 7.4, 20% glycerol, 2 mM EDTA, 1 mM DTT, 40 μM DEVD-AFC (55) (Enzo Biochem, Farmingdale, USA, #ALX260032M005), 5 μL of lysis buffer, and 5 μL of cellular lysate was added to each well. In control wells an extra 5 μL of lysis buffer was added in place of the cellular lysate. The plates were prepared on ice to limit enzymatic activity. The plates were read on a Bio-Tek Synergy Neo plate reader at an excitation wavelength of 400 nm and an emission wavelength of 500 nm. Reads were taken every 60 sec for 1 h and caspase3/7 activity is expressed as fold change in nM/AFC/mg/min.

### Colony formation

Cells were seeded into 6-well plates at a density of 1000 cells/well and were allowed to attach overnight. The next day the medium was changed, and the cells were treated as described in the figure legends. The growth medium was changed every 3 days. After 14 days the cells were fixed in 3.7% formaldehyde and the cells were stained with crystal violet. Images of the plates were acquired on a flat-bed scanner.

### Immunofluorescence staining

Cells were seeded into 8-well chamber slides and were treated as indicated in the figure legends. Cells were fixed with 3.7% formaldehyde for 15 min, and then blocked in 2% BSA PBS-Triton X-100 (0.1%) for 1 h at room temperature. Cells were incubated overnight at 4°C in primary antibody at a (1:1,000) dilution in blocking buffer. Cells were washed 3x in PBS then were incubated in the appropriate secondary antibody at a (1:1,000) dilution for 1 h at room temperature. Cells were washed 3x in PBS then were mounted in ProLong Gold Antifade + DAPI (ThermoFisher, #P36935). Slides were cured overnight at room temperature, and then transferred to 4°C. Slides were imaged on a Nikon TE2000-U fluorescence microscope at 20x magnification.

For all immunofluorescence experiments, images were blinded with an R script before quantification. We repeated all immunofluorescence experiments at least three times and typically analyzed 5-10 fields per biological replicate. In total we analyzed at least 200 cells per experimental group, but in most cases over 1000 cells per experimental group. For subcellular localization experiments, data are represented as a stacked bar graph wherein the fraction of cells that have predominantly nuclear, pan-cellular, or cytosolic localization is plotted as a fraction of the total cells. A cell was considered to have “cytosolic” localization if there was a clear nuclear exclusion. Inversely a cell was described as having “nuclear” localization if the staining intensity was appreciably higher than in the cytosol. If there was no apparent difference between the nuclear and cytosolic staining, then the cell was described as having “pan-cellular” distribution.

### RNA-Seq sample/data processing

Total cellular RNA was extracted from drug-treated M229R cells using the Qiagen (Hilden, Germany) RNeasy kit (#74104) with three biological replicates per cell line. All RNA samples had a RIN score > 8. Libraries were prepared using the Illumina TruSeq Stranded mRNA Library Preparation Kit, prepared libraries were quality controlled and quantified using a Qubit and Labchip Bioanalyzer. Libraries were pooled and run on a NovaSeq6000 instrument. Sequencing was performed by 2 × 150 bp paired-end read format. Base calling was done by Illumina RTA and converted to FASTQ using bcl2fastq software. Sequencing was performed at a depth of approximately 30 M reads/sample. Quality control was performed on the FASTQ files using FastQC v0.11.5, and reads were trimmed using Trimmomatic v0.33. Reads were mapped using HISAT2 v2.1.0 and analyzed using HTSeq v0.6.1. Differential gene expression was calculated using edgeR. Raw RNA-Seq reads and processed HTSeq read counts are available on GEO under GSE145990. When appropriate RNA-Seq data was upper quintile normalized prior to analysis.

### Datasets

Sources for the previously published RNA-Seq data used in this study are as follows. M229P/R and M238P/R RNA-Seq data was downloaded from GSE75313 (16). UACC62P/R RNA-Seq data was previously generated by our group and was deposited under GSE115938 (47). The PRISM drug response dataset was downloaded from the DepMap data download portal (depmap.org/portal/download).

### LISA

Epigenetic landscape in silico subtraction analysis (LISA) was run on lisa.cistrome.org (56). Gene lists were filtered to include only significantly differentially expressed genes (FDR < 0.01). Gene set 1 was filtered to include only upregulated genes, and gene set 2 was filtered to include only downregulated genes. Only the top 500 genes were used in each list. In cases where there were fewer than 500 differentially expressed genes, only the genes which had an FDR < 0.01 were included in the analysis. The ChIP-Seq output data was plotted as a scatter plot of enrichments in the upregulated vs downregulated gene sets.

### Connectivity map analysis

The top 200 upregulated/downregulated genes (FDR < 0.01) were analyzed to identify CMap Classes which have similar gene expression perturbation signatures on the online clue.io portal. In cases where there were fewer than 200 upregulated or downregulated genes with an FDR < 0.01, only genes which passed the FDR cutoff were included in the analysis.

### OCTAD Datasets and RNA-Sequence processing

We used the same pipeline to process RNA-Seq samples from public databases such as TCGA, TARGET, GTEx, and SRA and compiled them into one single dataset called OCTAD (57). Whenever possible, RNA-Seq samples used in this study were processed using the same pipeline to mitigate batch effects. In addition, RUVg (58) was used to remove unwanted variation, and weakly expressed genes were removed while computing differentially expressed genes. Normalized raw counts were used for DE analysis and TPM was used for other analyses. The clustering of these samples with melanoma samples compared to non-melanoma primary tumor samples demonstrates the feasibility of performing differential expression analysis between cell lines and tissue samples (Fig. S1).

### Disease signature creation

Gene expression data from BRAFi-resistant melanoma cell lines was compared with either 50 healthy normal skin samples from the GTEx database, or to BRAF^V600E^-mutant melanoma tumor samples to generate BRAFi-resistance gene expression signatures. We used edgeR to perform DE analysis (log_2_ fold change > 1, adjusted p-value < 0.001) (59). The detailed data processing and parameter selection were detailed in the OCTAD study (57). The enrichment of the genes in the BRAFi-resistance gene signatures was computed with _ss_GSEA (60). The association of enrichment scores for both of the signatures with patient survival was computed and visualized using the survminer package. Patient mutation status and survival data were retrieved from cBioPortal (61). EnrichR was used for pathway enrichment analysis (62).

### Drug prediction

The LINCS database containing gene expression profiles for compound-treated cells has been widely used for candidate drug prediction in our previous studies (23,63). The LINCS library is comprised of 476,251 signatures and 22,268 genes including 978 landmark genes. The 1,974 mapped drugs listed in the Repurposing Hub were considered in this study (64). To compute RGES scores, we first ranked genes based on their expression values in each drug signature. An enrichment score for each set of up- and down-regulated disease genes was computed separately using a Kolmogorov–Smirnov-like statistic, followed by the combination of scores from both sides. The score is based on the number of the genes (up or down-regulated) at either the top or bottom of a drug-gene list ranked by expression change after drug treatment. One compound might have multiple available expression profiles because they were tested in various cell lines, drug concentrations, treatment durations, or even different replicates, resulting in multiple RGES for one drug-disease prediction. We termed this score summarized RGES (sRGES). The computation of RGES and the summarization RGES were detailed elsewhere and recently implemented as a standalone R package (57). Compounds were filtered to include only compounds that had a sample size greater than 1 in the LINCS L1000 dataset and were filtered to exclude compounds that were anti-neoplastic or were previously studied in melanoma. A sRGES threshold of −0.3 was the cutoff for compounds which effectively reversed the BRAFi resistance signature.

## Results

### Identification of compounds which reverse a BRAFi resistance signature

We employed a systems-based approach to identify compounds that reverse an experimentally derived BRAFi resistance signature (Fig. 1A). This approach was originally proposed in the Connectivity Map project (21), and was extended in other studies (24,65), including a recent study from the Chen lab (23) which used sRGES to quantify the reversal potency and demonstrated its positive correlation with drug efficacy. Sample collection, signature creation, sRGES computation, and *in silico* validation were streamlined in the OCTAD pipeline which was described in the Materials and Methods section. This approach has been applied to identify potential therapeutic compounds for primary cancers, but this study is our first attempt to apply this method to study drug resistance.

**Figure 1.**
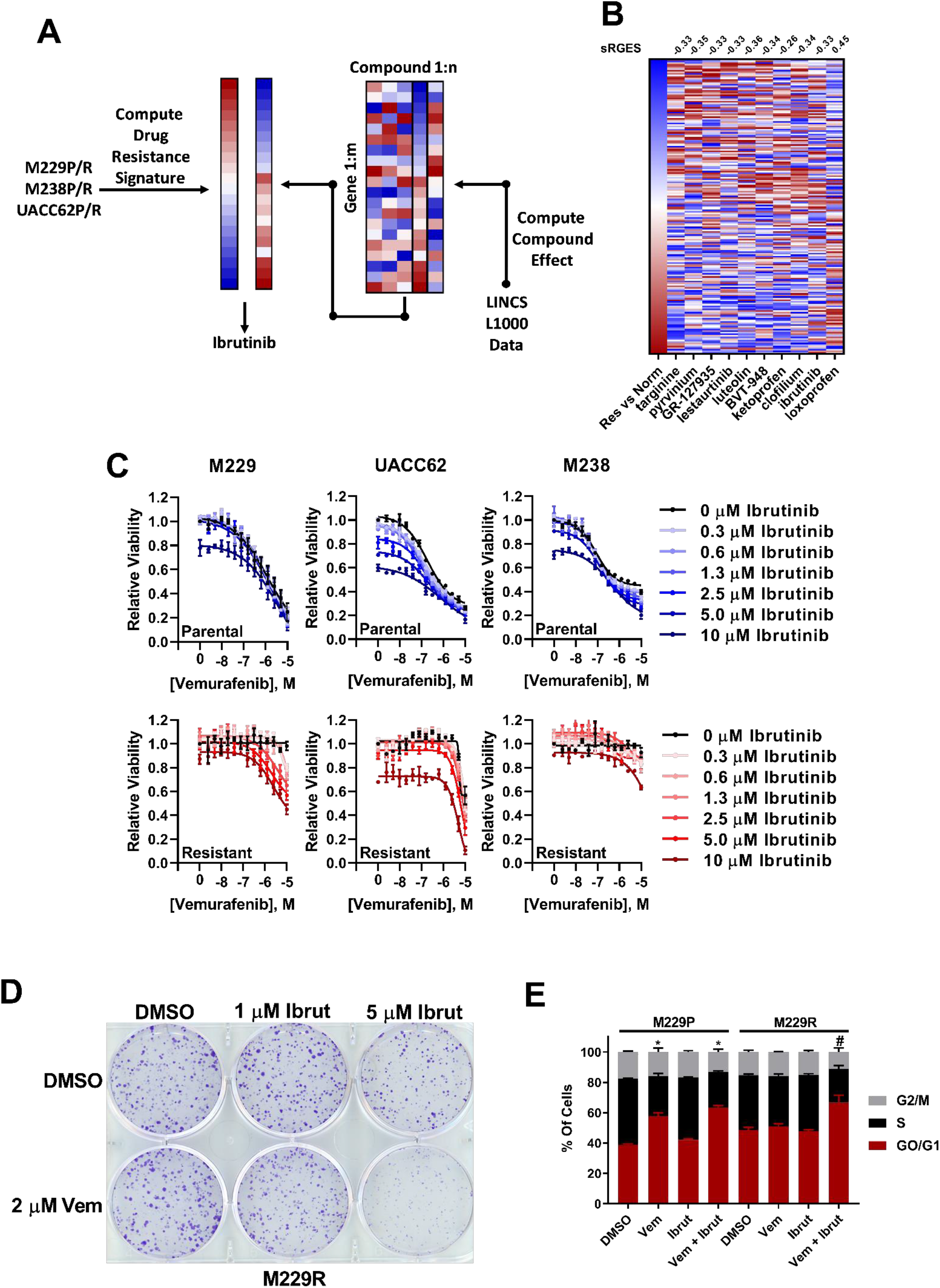
Ibrutinib re-sensitizes BRAFi-resistant cells to vemurafenib. **A.** Diagram of drug resistance and reversal signatures. **B.** The BRAFi-resistance signature was computed by comparing BRAFi-resistant cell lines and normal tissue samples. Red boxes indicate upregulated genes, and blue boxes indicate downregulated genes. Loxoprofen was included as a control, since this compound was not predicted to reverse the BRAFi-resistance signature. For compounds with multiple gene expression profiles, the profile with the median RGES was chosen for visualization. The sRGES values for the BRAFi-resistance signature and the compound-treated signatures are listed above the heatmap. **C.** M229P/R, UACC62P/R, and M238P/R cells were treated in a dose response matrix of ibrutinib (top concentration 10 μM, ½ dilution series) and vemurafenib (top concentration 10 μM, ½ dilution series). After 72 h, viability was measured with CellTiter-Glo. (n = 3 biological replicates) **D.** M229R cells were seeded into a 6-well plate at a density of 5,000 cells/well. The next day the cells were treated with the indicated concentrations of vemurafenib and ibrutinib. The colony formation assay was performed and analyzed as described in materials and methods. (n = 3 biological replicates) **E.** M229P/R cells were treated with −/+ 2 μM vemurafenib, −/+ 1 or 5 μM ibrutinib for 72 h. The cells were stained and analyzed by flow cytometry as described in materials and methods (n = 3 biological replicates). Significant differences of G0/G1 for compound treated samples vs the relevant DMSO control are indicated (One-way ANOVA, * p < 0.01 vs M229P-DMSO, # p < 0.01 vs M229R-DMSO).

We collected three datasets that include RNA-Seq profiles of parental and BRAFi-resistant melanoma cell lines (M229P/R, M238P/R, and UACC62P/R) with 2 biological replicates for each cell line. Initially we compared the profiles of parental and resistant cell lines for each dataset, but the gene signature did not effectively predict compound response using the CTRPv2 data; therefore, we decided to compare the resistant samples to healthy skin samples (n = 558) in the OCTAD database. We then used the most variable genes to select the 50 samples with the best correlation between healthy skin samples and BRAFi resistant samples. The comparison between these samples resulted in 191 DE genes that were included in the LINCS 978 landmark genes (log_2_ fold-change >1 and adjusted p-value < 0.001). The prediction identified 245 compounds with sRGES lower than −0.3. To computationally validate the predictions and tune parameters, we correlated the sRGES and compound sensitivity data for UACC62P cells in the CTRPv2 dataset (Fig. S2). The significant correlation (Spearman: 0.47, p-value: 1.6e-9) suggests that sRGES predictions are effective in predicting compound sensitivity in melanoma. Since one compound may be profiled against multiple cell lines in the LINCS L1000 dataset, we filtered RGES values by the mean score, standard deviation, and number of occurrences, and then performed enrichment analysis to confirm consistency across multiple cell lineages.

Nine compounds that reversed the BRAFi resistance gene expression signature *in silico* (Fig. 1B/Table S2) were selected and were examined for their ability to inhibit growth of matched parental and BRAFi-resistant melanoma cell lines. We identified 4 compounds that reduce cell viability in both M229P and M229R cells, with no apparent selectivity for one over the other (Fig. S3). This lack of selectivity is likely because both the parental and resistant cells were compared to normal tissue, instead of being directly compared against each other. Next, we created a gene expression resistance signature consisting of 87 genes by comparing the gene expression data from the resistant cell lines with BRAF^V600E^-mutant primary melanoma tumor samples in the OCTAD database. The expression signature is significantly associated with poor overall survival in melanoma patients with BRAF^V600E^ mutations (p = 0.006, Cox model), but not with BRAF^WT^ melanoma patients (p =0.028), suggesting that this gene expression signature may be clinically relevant (Fig. S4). With this new signature 3/9 of the compounds (ibrutinib, pyrvinium, and lestaurtinib) were among the top 5% of compounds identified, with ibrutinib being the most effective in reversing the BRAFi resistance signature (Fig. S5).

### Ibrutinib re-sensitizes BRAFi-resistant cells to vemurafenib

We reasoned that compounds which significantly reverse a BRAFi resistance gene expression signature should also reverse BRAFi resistance in melanoma cells in an experimental setting. To test this hypothesis, we profiled the synergy between vemurafenib and the top 9 hits from the computational screen in a 14×7 concentration response matrix with vemurafenib to identify compounds that can potentiate vemurafenib response. Out of the top 9 compounds identified in our screen, only ibrutinib reversed BRAFi resistance (Fig. 1C, red curves, Fig. S6). One interesting observation is that while the computational screen was performed using RNA-Seq data from all three isogenic parental and resistant cell line pairs, only M229R was re-sensitized to vemurafenib by ibrutinib. Synergistic growth inhibition was also observed in a long-term colony formation assay, which was more apparent with higher concentrations of ibrutinib (Fig. 1D). Since BRAF inhibitors arrest melanoma cells at the G1 checkpoint, if ibrutinib is truly re-sensitizing the resistant cells to vemurafenib it should also re-sensitize the cells to vemurafenib-induced G1 arrest. M229P cells accumulate in G0/G1 state during vemurafenib treatment but M229R cells do not. Consistent with re-sensitization we found that accumulation of M229R cells in G0/G1 is restored upon treatment with the combination of vemurafenib and ibrutinib (Fig. 1E). There was also an increased level of Annexin V-positive cells in the combination-treated group, although there was no change in Caspase 3/7 activity (Fig. S7). Taken together, these data suggest that ibrutinib re-sensitizes a subset of BRAFi-resistant cell lines to vemurafenib.

### BTK deletion or inhibition does not re-sensitize BRAFi-resistant cells to vemurafenib

Since ibrutinib is known to have targets other than BTK (37,66,67) we wanted to know whether BTK was responsible for BRAFi resistance. To test this hypothesis experimentally, we generated BTK knock out cell pools using CRISPR. BTK mRNA expression is low in both M229P and M229R (Fig. S8) cells making it impossible to assay knockout efficiency by immunoblotting, so we measured knockout efficiency by Sanger sequencing of gDNA amplicons which contain the region of the CRISPR cut site. The Sanger sequencing traces were subsequently de-convoluted with the TIDE algorithm (54) to identify the fraction of cells that had functional knockout (Fig. 2A/S9). Using this approach, we found that the functional knockout efficiency was approximately 70%. Even though ibrutinib is used clinically as a BTK inhibitor, deletion of BTK did not alter the vemurafenib response in either the parental or resistant cells (Fig. 2B). This suggested to us that ibrutinib may be re-sensitizing the cells through off-target inhibition of other kinases instead of by on-target inhibition of BTK. Since acalabrutinib is a BTK inhibitor analog of ibrutinib with significantly reduced off-target activity (66,67), we asked whether acalabrutinib reverses BRAFi resistance. Consistent with our hypothesis, acalabrutinib failed to re-sensitize BRAFi-resistant cells to vemurafenib (Fig. 2C). Taken together, these data show that the effect of ibrutinib to re-sensitize BRAFi-resistant cells to vemurafenib is independent of on-target BTK inhibition.

**Figure 2.**
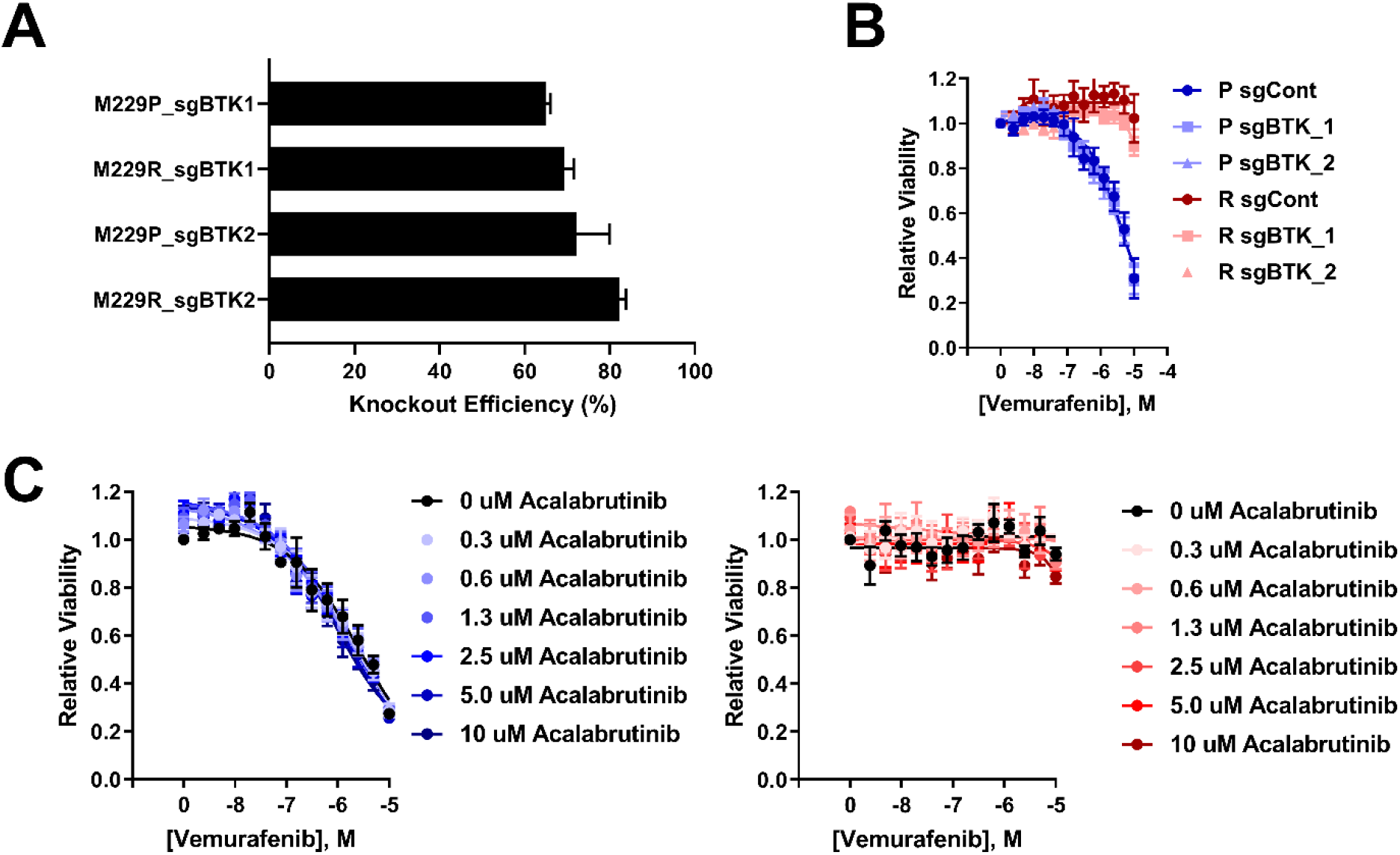
BTK deletion or inhibition does not alter vemurafenib sensitivity. **A.** M229P/R BTK^KO^ cells were generated as described in Materials and Methods. Sanger sequencing was performed to measure the extent of BTK deletion in M229P/R cell pools. The fraction of cells with functional BTK deletion was quantified with TIDE (n = 3 biological replicates)/ **B.** M229P/R sgControl and sgBTK cells were treated with 14 concentrations of vemurafenib (10 μM top concentration, ½ dilution series) and, after 72 h, viability was measured with CellTiter-Glo as described in Materials and Methods. (n = 3 biological replicates) **C.** M229P/R cells were treated with 7 different concentrations of acalabrutinib (10 μM top concentration, ½ dilution series) and 14 different concentrations of vemurafenib (10 μM top concentration, ½ dilution series). After 72 h, viability was measured with CellTiter-Glo. (n = 3 biological replicates)

### Transcriptional response to ibrutinib treatment

To better understand how ibrutinib re-sensitizes BRAFi-resistant cells to vemurafenib we performed RNA-seq on M229R cells after treatment with vemurafenib, ibrutinib, acalabrutinib, or combinations. Consistent with the observation that ibrutinib, but not acalabrutinib, re-sensitizes BRAFi-resistant cells to vemurafenib we found that there were 101 differentially expressed genes (FDR < 0.01) with ibrutinib treatment while there were no differentially expressed genes with acalabrutinib treatment (Fig. 3A). Compared to single agent treatment, there was a synergistic induction of differential gene expression with the combination of ibrutinib and vemurafenib (V+I) and V+I significantly reversed the BRAFi resistance signature used in the compound sensitivity predictions (Spearman correlation = −0.25, p-value = 0.0007) (Fig. 3B). We then identified networks of differentially expressed genes in cells cultured in the presence of ibrutinib or V+I. With either single agent ibrutinib or the combination of ibrutinib and vemurafenib, the gene networks were primarily associated with development of various organs (Fig. S10). To understand the effect of ibrutinib on melanoma cells in greater detail, we profiled transcriptional regulators that are predicted to be altered in cells cultured with ibrutinib or the combination of ibrutinib and vemurafenib using LISA (56) to identify transcription factors which may contribute to the differential gene expression in compound-treated cells. Among the top transcription regulators identified were YAP1 and two transcription factors, TEAD1 and TEAD4, which are bound by YAP1 (Fig. 3C). Interestingly, this enrichment was observed in genes that are both downregulated by ibrutinib treatment and genes that are upregulated by ibrutinib treatment. It is possible that this could be because YAP1 can function as a transcriptional repressor in addition to its canonical role as a transcriptional co-activator (68).

**Figure 3.**
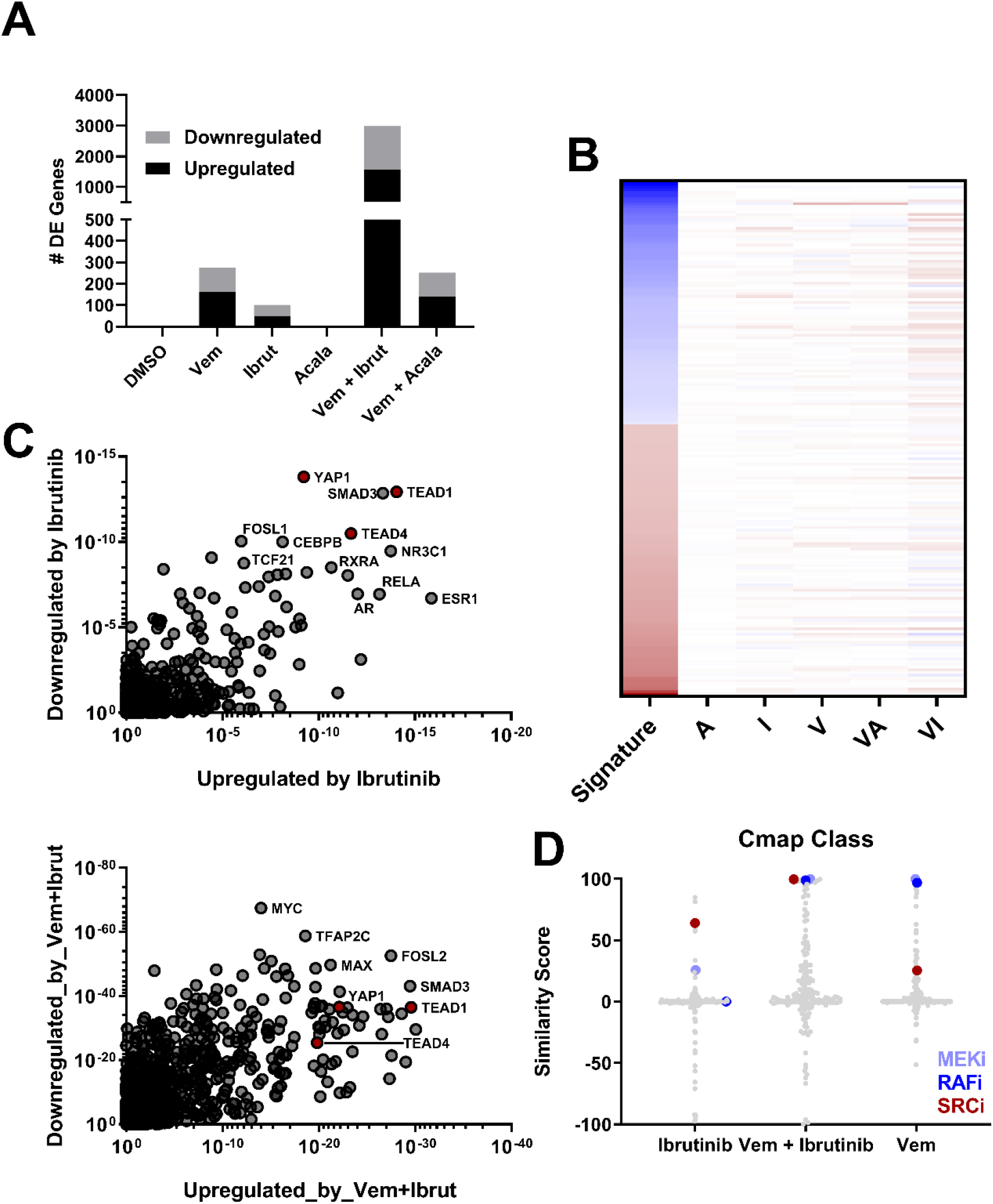
Transcriptional response to ibrutinib treatment. **A.** M229R cells were treated with DMSO, vemurafenib (2 μM), ibrutinib (5 μM), acalabrutinib (5 μM), or the combination of ibrutinib/acalabrutinib and vemurafenib. After 24 h RNA was extracted and RNA-Seq was performed as described in the materials and methods. **B.** Expression of genes in the BRAFi resistance signature which was used in the initial screen where ibrutinib was identified. For each compound the profiles of the three replicates were merged by taking the median expression value. For each treatment group the fold change in gene expression was compared to the DMSO control. Red boxes indicate that the gene is upregulated, and blue boxes indicate that the gene is downregulated. Only treatment with vemurafenib + ibrutinib significantly reversed the BRAFi resistance signature (Spearman correlation = −0.25, p-value = 0.0007). **C.** LISA analysis of differentially expressed genes in the ibrutinib and vemurafenib + ibrutinib treatment groups. Data analysis was performed as described in Materials and Methods. X- and Y-axis values are enrichment p-values. **D.** CMap class analysis was performed as described in Materials and Methods. Transcriptional signatures of ibrutinib, vemurafenib, or vemurafenib + ibrutinib were compared to transcriptional signatures in the Cmap dataset.

We reasoned that inhibitors with the same functional target as ibrutinib should have a similar transcriptional signature to ibrutinib. To address this, we compared the gene expression signatures of ibrutinib- and vemurafenib-treated cells to the signatures of other compounds in the Connectivity Map (Cmap) dataset. SRC inhibitors had a highly similar transcriptional signature to that of ibrutinib (Fig. 3D). This observation is interesting since ibrutinib, but not acalabrutinib, has significant off-target activity against multiple SRC family kinases (SFKs) (Fig. S11) (66,67). Another interesting observation was that the transcriptional signature of aromatase inhibitors was similar to that of ibrutinib, especially since expression of androgen receptor target genes was significantly enriched (Fig. 3C). However, treatment with several aromatase inhibitors did not alter BRAFi response in M229R cells (Fig. S12) suggesting that ibrutinib does not affect BRAFi sensitivity by modulating aromatase activity. As a further support that the method that we employed here works, we also performed the same comparison with vemurafenib-treated cells and found high similarity with BRAF and MEK inhibitors in the Cmap dataset, which is consistent with the pharmacology of vemurafenib. Together, these results suggest that ibrutinib alters YAP1 activity and the effects of ibrutinib on melanoma cells may be due to off-target anti-SFK activity.

### Ibrutinib reduces the nuclear accumulation of YAP1

YAP1 has been previously implicated in BRAFi resistance (4,44–47), so it is critical to understand whether ibrutinib is altering YAP1 activity. Transcriptionally inactive YAP1 is sequestered in the cytosol and upon various stimuli YAP1 can translocate into the nucleus where it modulates gene transcription. As we previously demonstrated (47), M229R cells have an increased nuclear/cytosolic ratio of YAP1 localization. Consistent with our computational predictions, ibrutinib reduced the proportion of cells with nuclear YAP1 localization; acalabrutinib did not have any effect on YAP1 localization (Fig. 4A/B). Interestingly, ibrutinib did not have any effect on YAP1 localization in M238R or UACC62R cells despite the fact that both resistant lines had elevated levels of nuclear YAP1 (Fig. 4C/D). It is possible that YAP1 could be regulated through other mechanisms in these cells, perhaps by control of serine phosphorylation by MST1/LATS (39). These data are consistent with our observation that ibrutinib re-sensitizes M229R cells to vemurafenib but only has a minor effect on M238R and UACC62R cells. We also observed an increase in the fraction of cells with predominantly nuclear TAZ localization in all three cell lines but neither ibrutinib not acalabrutinib altered TAZ localization (Fig. S13). Taken together, these data suggest that in a subset of BRAFi-resistant melanoma ibrutinib can alter YAP1 activity, which may contribute to re-sensitization to BRAFi treatment.

**Figure 4.**
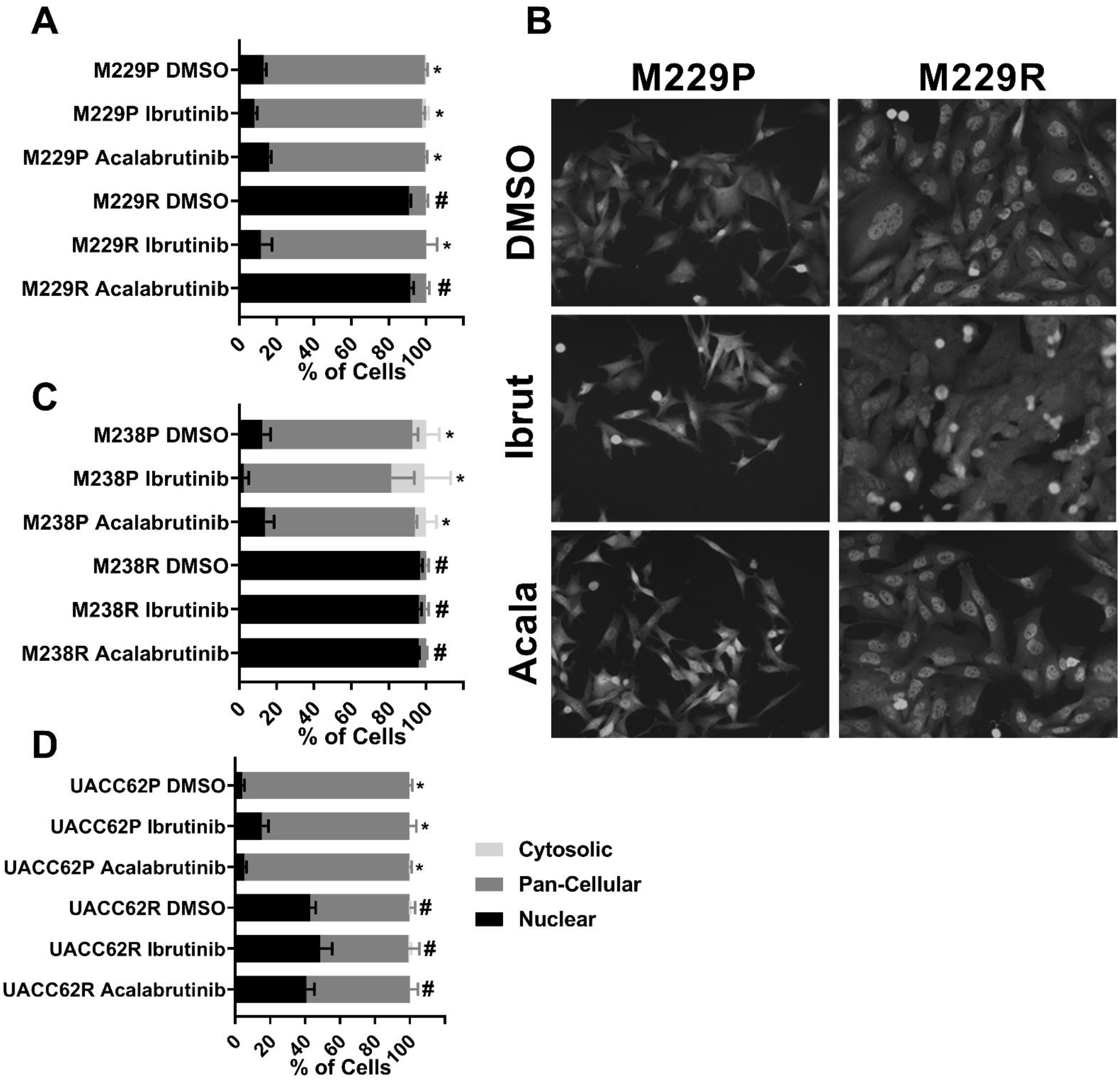
Ibrutinib blocks YAP1 nuclear localization. **A.** M229P/R cells were stained with an anti-YAP1 antibody as described in the materials and methods section. The percentage of cells with nuclear, cytosolic, or pan-cellular YAP1 localization was quantified as described in the materials and methods section. **B.** Representative images from the experiment in Fig 4A. **C.** M238P/R or **D.** UACC62P/R cells were stained with an anti-YAP1 antibody as described in the materials and methods section. The percentage of cells with nuclear, cytosolic, or pan-cellular YAP1 localization was quantified as described in the materials and methods section. Statistical analysis (one-way ANOVA) was performed on % of cells with nuclear YAP1 localization where p < 0.01 was considered statistically significant. Bars marked with # indicate a statistically significant difference when compared with DMSO-treated parental cells and bars marked with * indicate a statistically significant difference when compared with DMSO-treated resistant cells (n = 3 biological replicates for all imaging experiments).

## Discussion

In this study we used an unbiased computational approach to identify compounds that reverse a gene expression signature for BRAFi resistance. This identified a role for ibrutinib in re-sensitizing a subset of melanoma cells with acquired BRAFi resistance to vemurafenib. Our data show that this is not due to on-target BTK inhibition, but is due to off-target inhibition, presumably of at least one SFK. Other studies have also identified a role for SFKs in BRAFi resistance (69–71), further supporting the idea that off-target anti-SFK activity of potential melanoma therapeutics may be mechanistically important. One study in particular characterized a novel dual RAF/SRC inhibitor which retains activity against melanoma tumors which had previously developed resistance to dabrafenib/trametinib therapy (72).

Additionally, we found that ibrutinib, but not acalabrutinib, prevents the nuclear accumulation of YAP1, which would render YAP1 transcriptionally inactive (73). There is evidence in the literature that SFK inhibition may be critical in modulation of YAP1 activation. For example, YES1, a SFK that is bound by ibrutinib (37), phosphorylates and activates YAP1 (43). Other SFKs including LCK, as well as SRC itself, have also been demonstrated to modulate YAP1 activation (74,75), suggesting that modulation of YAP1 activity could be a general feature of SFKs.

Understanding the polypharmacology of ibrutinib will be critical for effectively re-purposing ibrutinib, an FDA approved drug, or related SFK inhibitors for the treatment of BRAFi-resistant melanoma. There is currently an ongoing clinical trial testing ibrutinib as a single agent therapy in patients with treatment-refractory metastatic melanoma (NCT02581930). Based on our findings that ibrutinib treatment alone is ineffective in BRAFi resistant or BRAFi-naïve (Fig. S14) melanoma cell lines, we would not expect a significant efficacy of ibrutinib as a single agent in the clinic. However, our data do suggest ibrutinib may re-sensitize a subset of resistant melanoma to BRAF inhibitors. Beyond melanoma, ibrutinib is used clinically to treat mantle cell lymphoma and chronic lymphocytic leukemia, and YAP1 has been implicated in both diseases (76,77). So the findings from this study may also be important in understanding differences in treatment response in these patients as well. It would be important to identify biomarkers, perhaps related to YAP1 activity or nuclear localization that would predict activity of SFK inhibition in BRAFi-resistant melanomas.

In this study we demonstrate that ibrutinib re-sensitizes a subset of BRAFi-resistant melanoma cells to vemurafenib. Mechanistically, we propose a model in which off-target SFK inhibition results in decreased YAP1 activity. The translational potential of this research is increased by the fact that ibrutinib is already FDA-approved, and thus can be used off-label for other indications. These data would suggest that ibrutinib or other SFK inhibitors, many of which are already FDA-approved, could have utility in BRAFi/MEKi-resistant melanoma tumors, as well as other YAP1-driven cancers.

## Bibliography

1. Hodis E, Watson IR, Kryukov GV, Arold ST, Imielinski M, Theurillat JP, et al. A landscape of driver mutations in melanoma. Cell 2012;150:251–63

2. Larkin J, Ascierto PA, Dreno B, Atkinson V, Liszkay G, Maio M, et al. Combined vemurafenib and cobimetinib in BRAF-mutated melanoma. N Engl J Med 2014;371:1867–76

3. Robert C, Karaszewska B, Schachter J, Rutkowski P, Mackiewicz A, Stroiakovski D, et al. Improved overall survival in melanoma with combined dabrafenib and trametinib. N Engl J Med 2015;372:30–9

4. Hugo W, Shi H, Sun L, Piva M, Song C, Kong X, et al. Non-genomic and Immune Evolution of Melanoma Acquiring MAPKi Resistance. Cell 2015;162:1271–85

5. Johnson DB, Menzies AM, Zimmer L, Eroglu Z, Ye F, Zhao S, et al. Acquired BRAF inhibitor resistance: A multicenter meta-analysis of the spectrum and frequencies, clinical behaviour, and phenotypic associations of resistance mechanisms. Eur J Cancer 2015;51:2792–9

6. Van Allen EM, Wagle N, Sucker A, Treacy DJ, Johannessen CM, Goetz EM, et al. The genetic landscape of clinical resistance to RAF inhibition in metastatic melanoma. Cancer Discov 2014;4:94–109

7. Shi H, Moriceau G, Kong X, Lee MK, Lee H, Koya RC, et al. Melanoma whole-exome sequencing identifies (V600E)B-RAF amplification-mediated acquired B-RAF inhibitor resistance. Nat Commun 2012;3:724

8. Shi H, Hugo W, Kong X, Hong A, Koya RC, Moriceau G, et al. Acquired resistance and clonal evolution in melanoma during BRAF inhibitor therapy. Cancer Discov 2014;4:80–93

9. Poulikakos PI, Persaud Y, Janakiraman M, Kong X, Ng C, Moriceau G, et al. RAF inhibitor resistance is mediated by dimerization of aberrantly spliced BRAF(V600E). Nature 2011;480:387–90

10. Emery CM, Vijayendran KG, Zipser MC, Sawyer AM, Niu L, Kim JJ, et al. MEK1 mutations confer resistance to MEK and B-RAF inhibition. Proc Natl Acad Sci U S A 2009;106:20411–6

11. Nazarian R, Shi H, Wang Q, Kong X, Koya RC, Lee H, et al. Melanomas acquire resistance to B-RAF(V600E) inhibition by RTK or N-RAS upregulation. Nature 2010;468:973–7

12. Whittaker SR, Theurillat JP, Van Allen E, Wagle N, Hsiao J, Cowley GS, et al. A genome-scale RNA interference screen implicates NF1 loss in resistance to RAF inhibition. Cancer Discov 2013;3:350–62

13. Johannessen CM, Boehm JS, Kim SY, Thomas SR, Wardwell L, Johnson LA, et al. COT drives resistance to RAF inhibition through MAP kinase pathway reactivation. Nature 2010;468:968–72

14. Saei A, Palafox M, Benoukraf T, Kumari N, Jaynes PW, Iyengar PV, et al. Loss of USP28-mediated BRAF degradation drives resistance to RAF cancer therapies. J Exp Med 2018;215:1913–28

15. Shen CH, Kim SH, Trousil S, Frederick DT, Piris A, Yuan P, et al. Loss of cohesin complex components STAG2 or STAG3 confers resistance to BRAF inhibition in melanoma. Nat Med 2016;22:1056–61

16. Song C, Piva M, Sun L, Hong A, Moriceau G, Kong X, et al. Recurrent Tumor Cell-Intrinsic and-Extrinsic Alterations during MAPKi-Induced Melanoma Regression and Early Adaptation. Cancer Discov 2017;7:1248–65

17. Shaffer SM, Dunagin MC, Torborg SR, Torre EA, Emert B, Krepler C, et al. Rare cell variability and drug-induced reprogramming as a mode of cancer drug resistance. Nature 2017;546:431–5

18. Das Thakur M, Salangsang F, Landman AS, Sellers WR, Pryer NK, Levesque MP, et al. Modelling vemurafenib resistance in melanoma reveals a strategy to forestall drug resistance. Nature 2013;494:251–5

19. Hong A, Moriceau G, Sun L, Lomeli S, Piva M, Damoiseaux R, et al. Exploiting Drug Addiction Mechanisms to Select against MAPKi-Resistant Melanoma. Cancer Discov 2018;8:74–93

20. Moriceau G, Hugo W, Hong A, Shi H, Kong X, Yu CC, et al. Tunable-combinatorial mechanisms of acquired resistance limit the efficacy of BRAF/MEK cotargeting but result in melanoma drug addiction. Cancer Cell 2015;27:240–56

21. Lamb J, Crawford ED, Peck D, Modell JW, Blat IC, Wrobel MJ, et al. The Connectivity Map: using gene-expression signatures to connect small molecules, genes, and disease. Science 2006;313:1929–35

22. Subramanian A, Narayan R, Corsello SM, Peck DD, Natoli TE, Lu X, et al. A Next Generation Connectivity Map: L1000 Platform and the First 1,000,000 Profiles. Cell 2017;171:1437–52 e17

23. Chen B, Ma L, Paik H, Sirota M, Wei W, Chua MS, et al. Reversal of cancer gene expression correlates with drug efficacy and reveals therapeutic targets. Nat Commun 2017;8:16022

24. Chen B, Wei W, Ma L, Yang B, Gill RM, Chua MS, et al. Computational Discovery of Niclosamide Ethanolamine, a Repurposed Drug Candidate That Reduces Growth of Hepatocellular Carcinoma Cells In Vitro and in Mice by Inhibiting Cell Division Cycle 37 Signaling. Gastroenterology 2017;152:2022–36

25. Jahchan NS, Dudley JT, Mazur PK, Flores N, Yang D, Palmerton A, et al. A drug repositioning approach identifies tricyclic antidepressants as inhibitors of small cell lung cancer and other neuroendocrine tumors. Cancer Discov 2013;3:1364–77

26. Antolin AA, Workman P, Mestres J, Al-Lazikani B. Polypharmacology in Precision Oncology: Current Applications and Future Prospects. Curr Pharm Des 2016;22:6935–45

27. Buchdunger E, Zimmermann J, Mett H, Meyer T, Muller M, Druker BJ, et al. Inhibition of the Abl protein-tyrosine kinase in vitro and in vivo by a 2-phenylaminopyrimidine derivative. Cancer Res 1996;56:100–4

28. Deininger MW, Goldman JM, Lydon N, Melo JV. The tyrosine kinase inhibitor CGP57148B selectively inhibits the growth of BCR-ABL-positive cells. Blood 1997;90:3691–8

29. Druker BJ, Tamura S, Buchdunger E, Ohno S, Segal GM, Fanning S, et al. Effects of a selective inhibitor of the Abl tyrosine kinase on the growth of Bcr-Abl positive cells. Nat Med 1996;2:561–6

30. Rubin BP, Schuetze SM, Eary JF, Norwood TH, Mirza S, Conrad EU, et al. Molecular targeting of platelet-derived growth factor B by imatinib mesylate in a patient with metastatic dermatofibrosarcoma protuberans. J Clin Oncol 2002;20:3586–91

31. Sjoblom T, Shimizu A, O’Brien KP, Pietras K, Dal Cin P, Buchdunger E, et al. Growth inhibition of dermatofibrosarcoma protuberans tumors by the platelet-derived growth factor receptor antagonist STI571 through induction of apoptosis. Cancer Res 2001;61:5778–83

32. Shimizu A, O’Brien KP, Sjoblom T, Pietras K, Buchdunger E, Collins VP, et al. The dermatofibrosarcoma protuberans-associated collagen type Ialpha1/platelet-derived growth factor (PDGF) B-chain fusion gene generates a transforming protein that is processed to functional PDGF-BB. Cancer Res 1999;59:3719–23

33. Greco A, Roccato E, Miranda C, Cleris L, Formelli F, Pierotti MA. Growth-inhibitory effect of STI571 on cells transformed by the COL1A1/PDGFB rearrangement. Int J Cancer 2001;92:354–60

34. Joensuu H, Roberts PJ, Sarlomo-Rikala M, Andersson LC, Tervahartiala P, Tuveson D, et al. Effect of the tyrosine kinase inhibitor STI571 in a patient with a metastatic gastrointestinal stromal tumor. N Engl J Med 2001;344:1052–6

35. Tuveson DA, Willis NA, Jacks T, Griffin JD, Singer S, Fletcher CD, et al. STI571 inactivation of the gastrointestinal stromal tumor c-KIT oncoprotein: biological and clinical implications. Oncogene 2001;20:5054–8

36. Lin A, Giuliano CJ, Palladino A, John KM, Abramowicz C, Yuan ML, et al. Off-target toxicity is a common mechanism of action of cancer drugs undergoing clinical trials. Sci Transl Med 2019;11

37. Klaeger S, Heinzlmeir S, Wilhelm M, Polzer H, Vick B, Koenig PA, et al. The target landscape of clinical kinase drugs. Science 2017;358

38. Zhao B, Wei X, Li W, Udan RS, Yang Q, Kim J, et al. Inactivation of YAP oncoprotein by the Hippo pathway is involved in cell contact inhibition and tissue growth control. Genes Dev 2007;21:2747–61

39. Oka T, Mazack V, Sudol M. Mst2 and Lats kinases regulate apoptotic function of Yes kinase-associated protein (YAP). J Biol Chem 2008;283:27534–46

40. Zhao B, Li L, Wang L, Wang CY, Yu J, Guan KL. Cell detachment activates the Hippo pathway via cytoskeleton reorganization to induce anoikis. Genes Dev 2012;26:54–68

41. Wada K, Itoga K, Okano T, Yonemura S, Sasaki H. Hippo pathway regulation by cell morphology and stress fibers. Development 2011;138:3907–14

42. Zhao B, Li L, Tumaneng K, Wang CY, Guan KL. A coordinated phosphorylation by Lats and CK1 regulates YAP stability through SCF(beta-TRCP). Genes Dev 2010;24:72–85

43. Rosenbluh J, Nijhawan D, Cox AG, Li X, Neal JT, Schafer EJ, et al. beta-Catenin-driven cancers require a YAP1 transcriptional complex for survival and tumorigenesis. Cell 2012;151:1457–73

44. Lin L, Sabnis AJ, Chan E, Olivas V, Cade L, Pazarentzos E, et al. The Hippo effector YAP promotes resistance to RAF- and MEK-targeted cancer therapies. Nat Genet 2015;47:250–6

45. Fisher ML, Grun D, Adhikary G, Xu W, Eckert RL. Inhibition of YAP function overcomes BRAF inhibitor resistance in melanoma cancer stem cells. Oncotarget 2017;8:110257–72

46. Kim MH, Kim J, Hong H, Lee SH, Lee JK, Jung E, et al. Actin remodeling confers BRAF inhibitor resistance to melanoma cells through YAP/TAZ activation. EMBO J 2016;35:462–78

47. Misek SA, Appleton KM, Dexheimer TS, Lisabeth EM, Lo RS, Larsen SD, et al. Rho-mediated signaling promotes BRAF inhibitor resistance in de-differentiated melanoma cells. Oncogene 2020;39:1466–83

48. Chen D, Sun Y, Wei Y, Zhang P, Rezaeian AH, Teruya-Feldstein J, et al. LIFR is a breast cancer metastasis suppressor upstream of the Hippo-YAP pathway and a prognostic marker. Nat Med 2012;18:1511–7

49. Orr BA, Bai H, Odia Y, Jain D, Anders RA, Eberhart CG. Yes-associated protein 1 is widely expressed in human brain tumors and promotes glioblastoma growth. J Neuropathol Exp Neurol 2011;70:568–77

50. Zhang W, Nandakumar N, Shi Y, Manzano M, Smith A, Graham G, et al. Downstream of mutant KRAS, the transcription regulator YAP is essential for neoplastic progression to pancreatic ductal adenocarcinoma. Sci Signal 2014;7:ra42

51. Tschaharganeh DF, Chen X, Latzko P, Malz M, Gaida MM, Felix K, et al. Yes-associated protein up-regulates Jagged-1 and activates the Notch pathway in human hepatocellular carcinoma. Gastroenterology 2013;144:1530–42 e12

52. Chaib I, Karachaliou N, Pilotto S, Codony Servat J, Cai X, Li X, et al. Co-activation of STAT3 and YES-Associated Protein 1 (YAP1) Pathway in EGFR-Mutant NSCLC. J Natl Cancer Inst 2017;109

53. Lui JW, Xiao S, Ogomori K, Hammarstedt JE, Little EC, Lang D. The Efficiency of Verteporfin as a Therapeutic Option in Pre-Clinical Models of Melanoma. J Cancer 2019;10:1–10

54. Brinkman EK, Chen T, Amendola M, van Steensel B. Easy quantitative assessment of genome editing by sequence trace decomposition. Nucleic Acids Res 2014;42:e168

55. Arango D, Parihar A, Villamena FA, Wang L, Freitas MA, Grotewold E, et al. Apigenin induces DNA damage through the PKCdelta-dependent activation of ATM and H2AX causing down-regulation of genes involved in cell cycle control and DNA repair. Biochem Pharmacol 2012;84:1571–80

56. Qin Q, Fan J, Zheng R, Wan C, Mei S, Wu Q, et al. Lisa: inferring transcriptional regulators through integrative modeling of public chromatin accessibility and ChIP-seq data. Genome Biol 2020;21:32

57. Zeng B, Glicksberg BS, Newbury P, Xing J, Liu K, Wen A, et al. OCTAD: an open workplace for virtually screening therapeutics targeting precise cancer patient groups using gene expression features. bioRxiv 2019:821546

58. Risso D, Ngai J, Speed TP, Dudoit S. Normalization of RNA-seq data using factor analysis of control genes or samples. Nat Biotechnol 2014;32:896–902

59. Robinson MD, McCarthy DJ, Smyth GK. edgeR: a Bioconductor package for differential expression analysis of digital gene expression data. Bioinformatics 2010;26:139–40

60. Hanzelmann S, Castelo R, Guinney J. GSVA: gene set variation analysis for microarray and RNA-seq data. BMC Bioinformatics 2013;14:7

61. Gao J, Aksoy BA, Dogrusoz U, Dresdner G, Gross B, Sumer SO, et al. Integrative analysis of complex cancer genomics and clinical profiles using the cBioPortal. Sci Signal 2013;6:pl1

62. Kuleshov MV, Jones MR, Rouillard AD, Fernandez NF, Duan Q, Wang Z, et al. Enrichr: a comprehensive gene set enrichment analysis web server 2016 update. Nucleic Acids Res 2016;44:W90–7

63. Chen B, Greenside P, Paik H, Sirota M, Hadley D, Butte AJ. Relating Chemical Structure to Cellular Response: An Integrative Analysis of Gene Expression, Bioactivity, and Structural Data Across 11,000 Compounds. CPT Pharmacometrics Syst Pharmacol 2015;4:576–84

64. Corsello SM, Bittker JA, Liu Z, Gould J, McCarren P, Hirschman JE, et al. The Drug Repurposing Hub: a next-generation drug library and information resource. Nat Med 2017;23:405–8

65. Sirota M, Dudley JT, Kim J, Chiang AP, Morgan AA, Sweet-Cordero A, et al. Discovery and preclinical validation of drug indications using compendia of public gene expression data. Sci Transl Med 2011;3:96 ra77

66. Patel V, Balakrishnan K, Bibikova E, Ayres M, Keating MJ, Wierda WG, et al. Comparison of Acalabrutinib, A Selective Bruton Tyrosine Kinase Inhibitor, with Ibrutinib in Chronic Lymphocytic Leukemia Cells. Clin Cancer Res 2017;23:3734–43

67. Herman SEM, Montraveta A, Niemann CU, Mora-Jensen H, Gulrajani M, Krantz F, et al. The Bruton Tyrosine Kinase (BTK) Inhibitor Acalabrutinib Demonstrates Potent On-Target Effects and Efficacy in Two Mouse Models of Chronic Lymphocytic Leukemia. Clin Cancer Res 2017;23:2831–41

68. Kim M, Kim T, Johnson RL, Lim DS. Transcriptional co-repressor function of the hippo pathway transducers YAP and TAZ. Cell Rep 2015;11:270–82

69. Fallahi-Sichani M, Becker V, Izar B, Baker GJ, Lin JR, Boswell SA, et al. Adaptive resistance of melanoma cells to RAF inhibition via reversible induction of a slowly dividing de-differentiated state. Mol Syst Biol 2017;13:905

70. Feddersen CR, Schillo JL, Varzavand A, Vaughn HR, Wadsworth LS, Voigt AP, et al. Src-Dependent DBL Family Members Drive Resistance to Vemurafenib in Human Melanoma. Cancer Res 2019;79:5074–87

71. Girotti MR, Pedersen M, Sanchez-Laorden B, Viros A, Turajlic S, Niculescu-Duvaz D, et al. Inhibiting EGF receptor or SRC family kinase signaling overcomes BRAF inhibitor resistance in melanoma. Cancer Discov 2013;3:158–67

72. Girotti MR, Lopes F, Preece N, Niculescu-Duvaz D, Zambon A, Davies L, et al. Paradox-breaking RAF inhibitors that also target SRC are effective in drug-resistant BRAF mutant melanoma. Cancer Cell 2015;27:85–96

73. Basu S, Totty NF, Irwin MS, Sudol M, Downward J. Akt phosphorylates the Yes-associated protein, YAP, to induce interaction with 14-3-3 and attenuation of p73-mediated apoptosis. Mol Cell 2003;11:11–23

74. Sugihara T, Werneburg NW, Hernandez MC, Yang L, Kabashima A, Hirsova P, et al. YAP Tyrosine Phosphorylation and Nuclear Localization in Cholangiocarcinoma Cells Are Regulated by LCK and Independent of LATS Activity. Mol Cancer Res 2018;16:1556–67

75. Lamar JM, Xiao Y, Norton E, Jiang ZG, Gerhard GM, Kooner S, et al. SRC tyrosine kinase activates the YAP/TAZ axis and thereby drives tumor growth and metastasis. J Biol Chem 2018

76. Byrd JC, Furman RR, Coutre SE, Flinn IW, Burger JA, Blum KA, et al. Targeting BTK with ibrutinib in relapsed chronic lymphocytic leukemia. N Engl J Med 2013;369:32–42

77. Wang ML, Rule S, Martin P, Goy A, Auer R, Kahl BS, et al. Targeting BTK with ibrutinib in relapsed or refractory mantle-cell lymphoma. N Engl J Med 2013;369:507–16

