## Supplemental Figures for "Ibrutinib blocks YAP1 activation and reverses BRAFi resistance in melanoma cells"

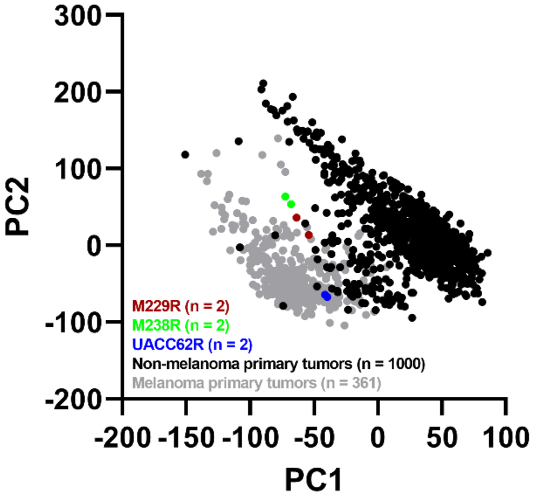


**Fig. S1: Principal Component Analysis of resistant cell line samples and tumor tissue samples.** Principal Component Analysis was performed on 361 primary melanoma tumors, 1,000 non-melanoma primary tumors, and BRAFi-resistant melanoma cell lines (n = 2 for 3 different cell lines).

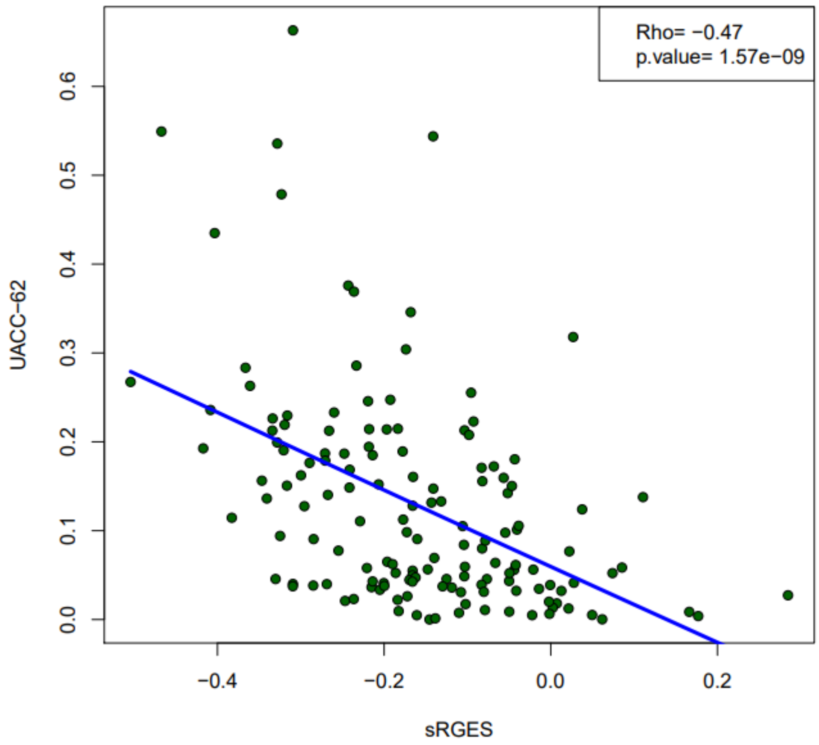


**Fig S2. Drug sensitivity correlates with sRGES drug response predictions.** Predicted drug sensitivity was calculated for UACC62P cells and was correlated with drug response data from the CTRPv2 dataset.

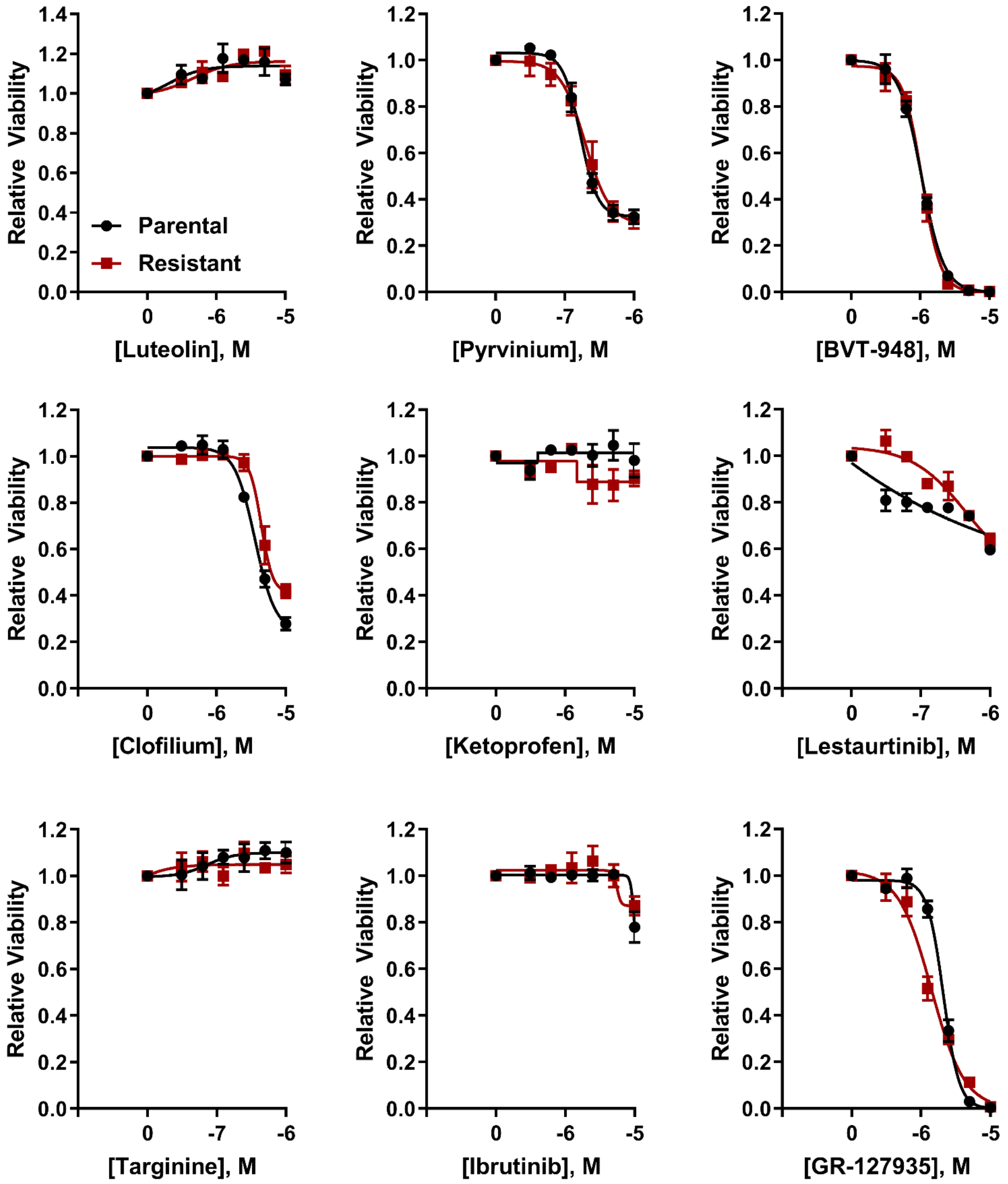


**Fig S3. Single-agent activity of compounds identified in the computational screen.** M229P (black lines) and M229R (red lines) cells were seeded into 384-well plates at a density of 1,000 cells/well. The next day the cells were treated with the indicated compounds. After 72 h viability was measured as described in materials and methods. The single agent response curves were derived from the experiment in Fig S2 but are re-plotted here as a separate figure to improve clarity and ease interpretation of the data.

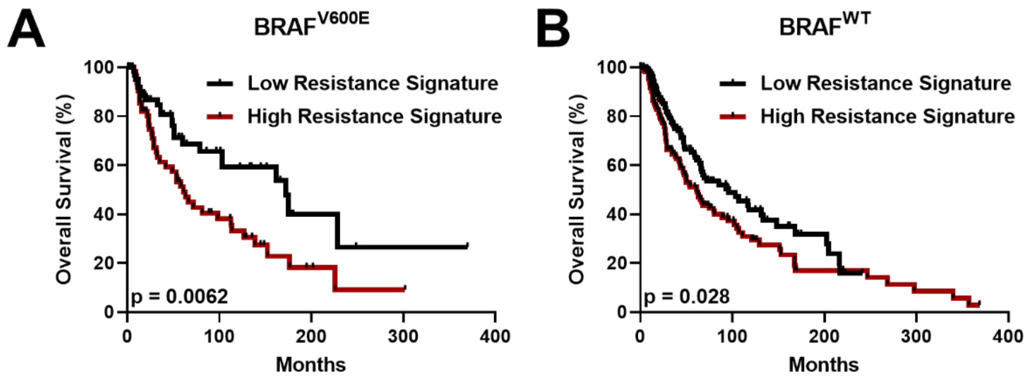


**Fig. S4. A BRAFi resistance signature is inversely correlated with melanoma overall survival.** The BRAFi-resistance gene expression signature was generated as described in the Materials and Methods section and expression of this signature was calculated for either **A.** BRAF$^{V600E}$ or **B.** BRAF$^{WT}$ melanoma tumors.

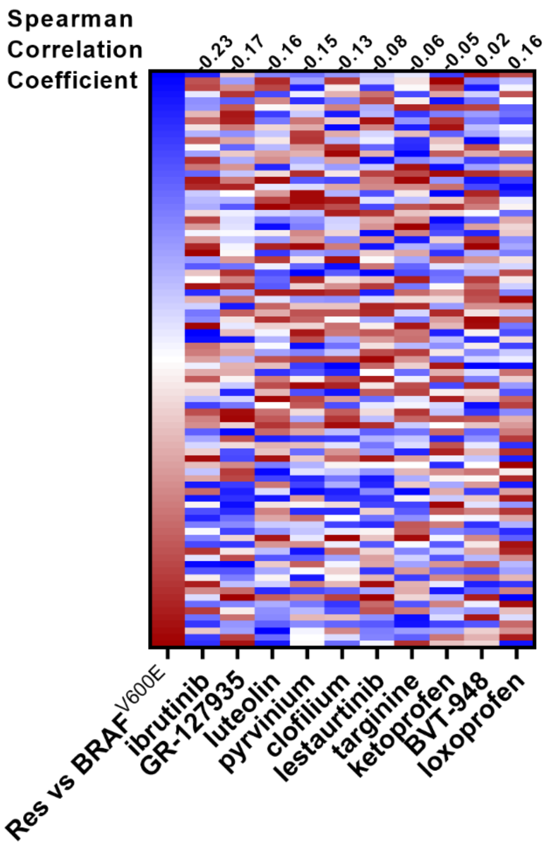


**Fig. S5. Identification of compounds that reverse a BRAFi resistance gene expression signature.** The resistance signature was computed by comparing resistance cell line samples and BRAF$^{V600E}$-mutant melanoma patient samples. Red boxes indicate that the gene is upregulated, and blue boxes indicate downregulated genes. Loxoprofen was included as a control since this compound was not predicted to reverse the BRAFi resistance signature. For compounds with multiple gene expression profiles, the profile with a median RGES was chosen for visualization. The correlation coefficients for the BRAFi-resistance signature and the compound-treated signatures are listed above the heatmap.

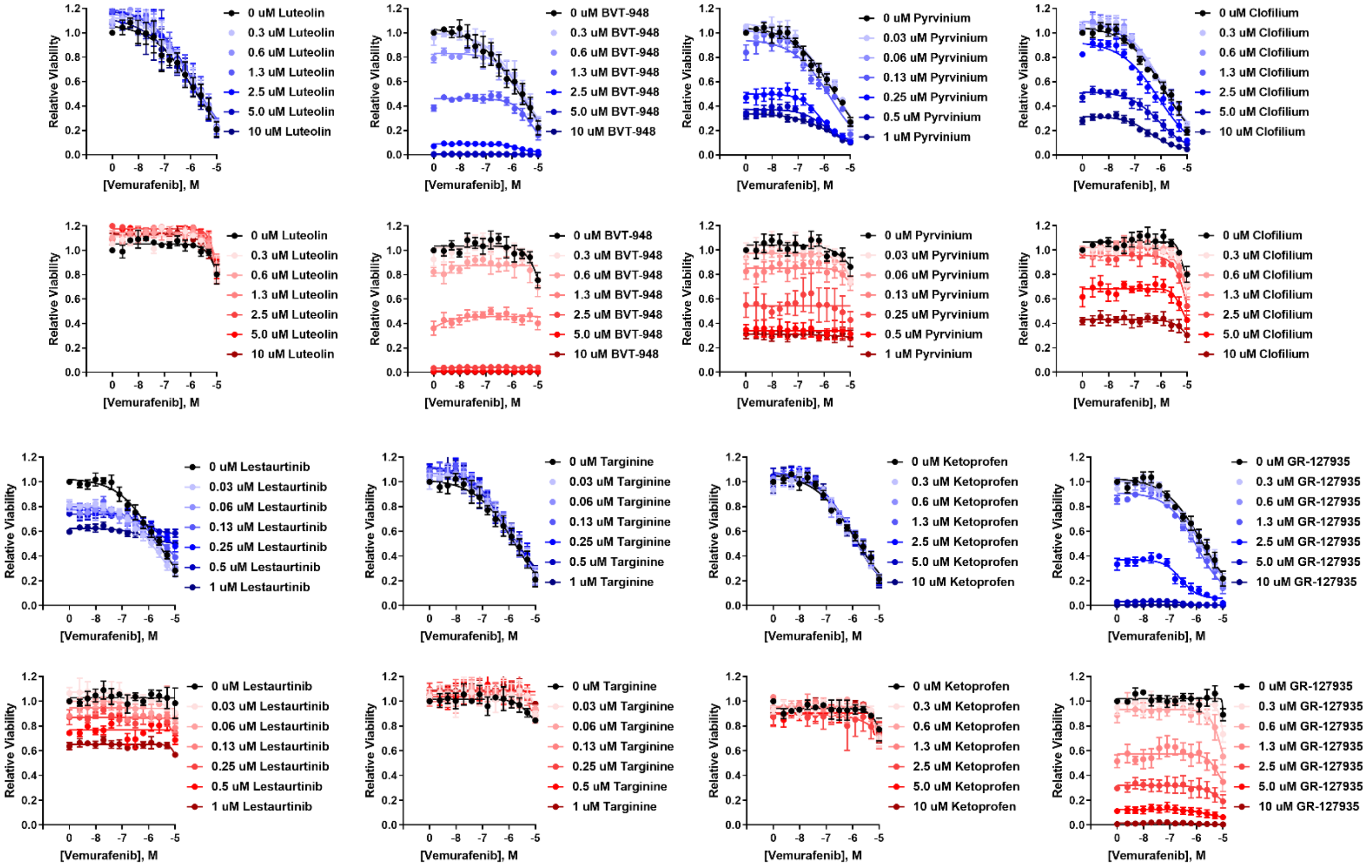


**Fig S6. Identification of compounds which re-sensitize BRAFi-resistant cells to vemurafenib.** M229P (blue lines) and M229R (red lines) cells were seeded into 384-well plates at a density of 1,000 cells/well. The next day the cells were treated with the indicated compounds. After 72 h viability was measured as described in materials and methods.

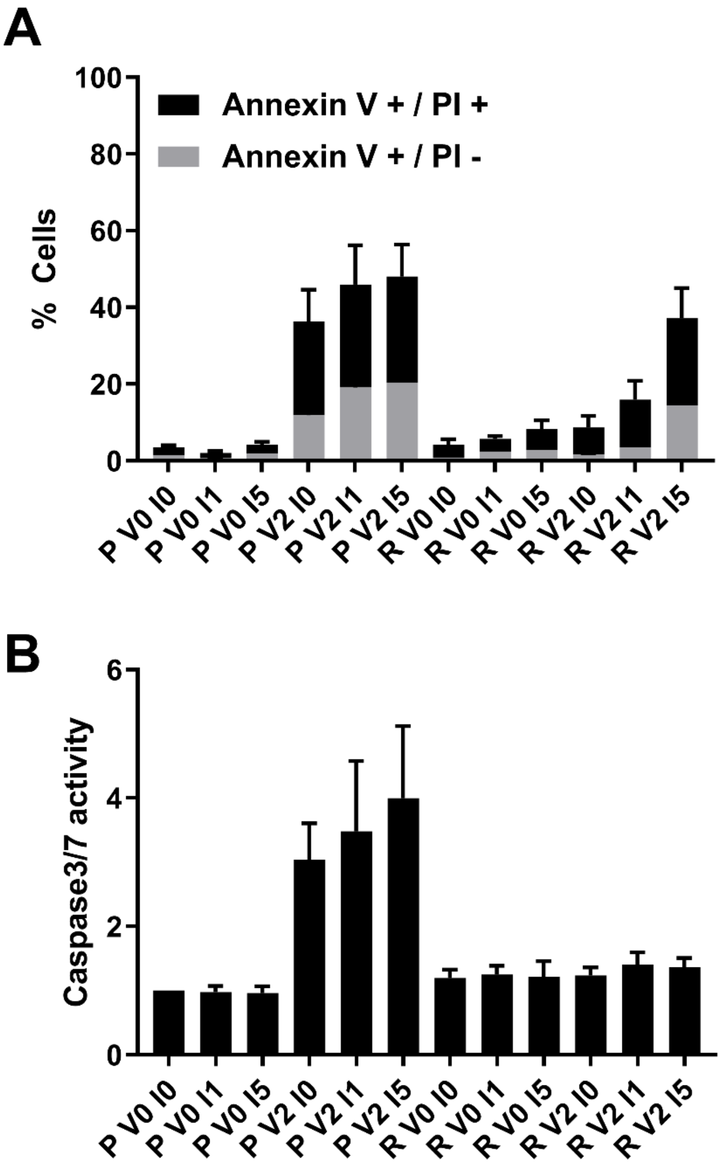


**Fig S7. The combination of vemurafenib and ibrutinib increases the number of Annexin V-positive cells but does not alter caspase3/7 activity. A.** The proportion of Annexin V and Propidium Iodide positive M229P/R cells was analyzed with flow cytometry as described in the Materials and Methods section. **B.** DEVD-AFC assays were used to evaluate caspase3/7 activity in M229P/R cellular lysates as described in the Materials and Methods section.

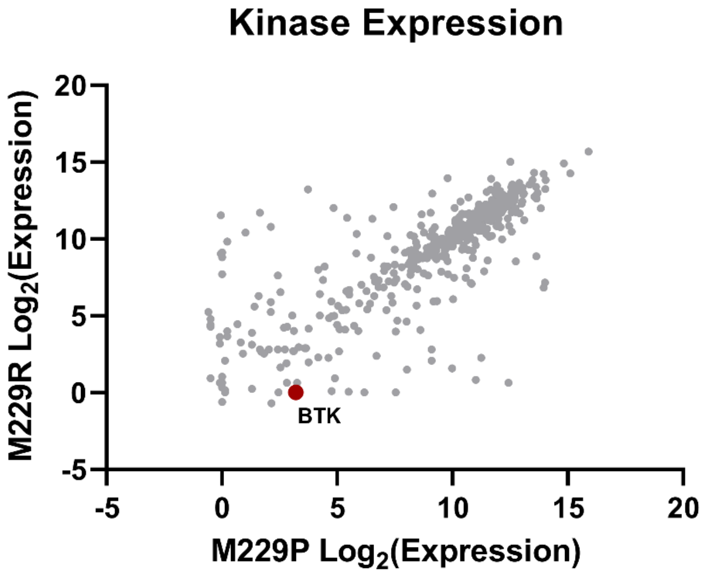


**Fig S8. BTK is weakly expressed in M229P/R cells.** RNA-Seq data for M229P/R cells was processed as described in Materials and Methods and expression of all protein kinases was compared. Relative to other kinases, the number of detected reads for BTK was low.

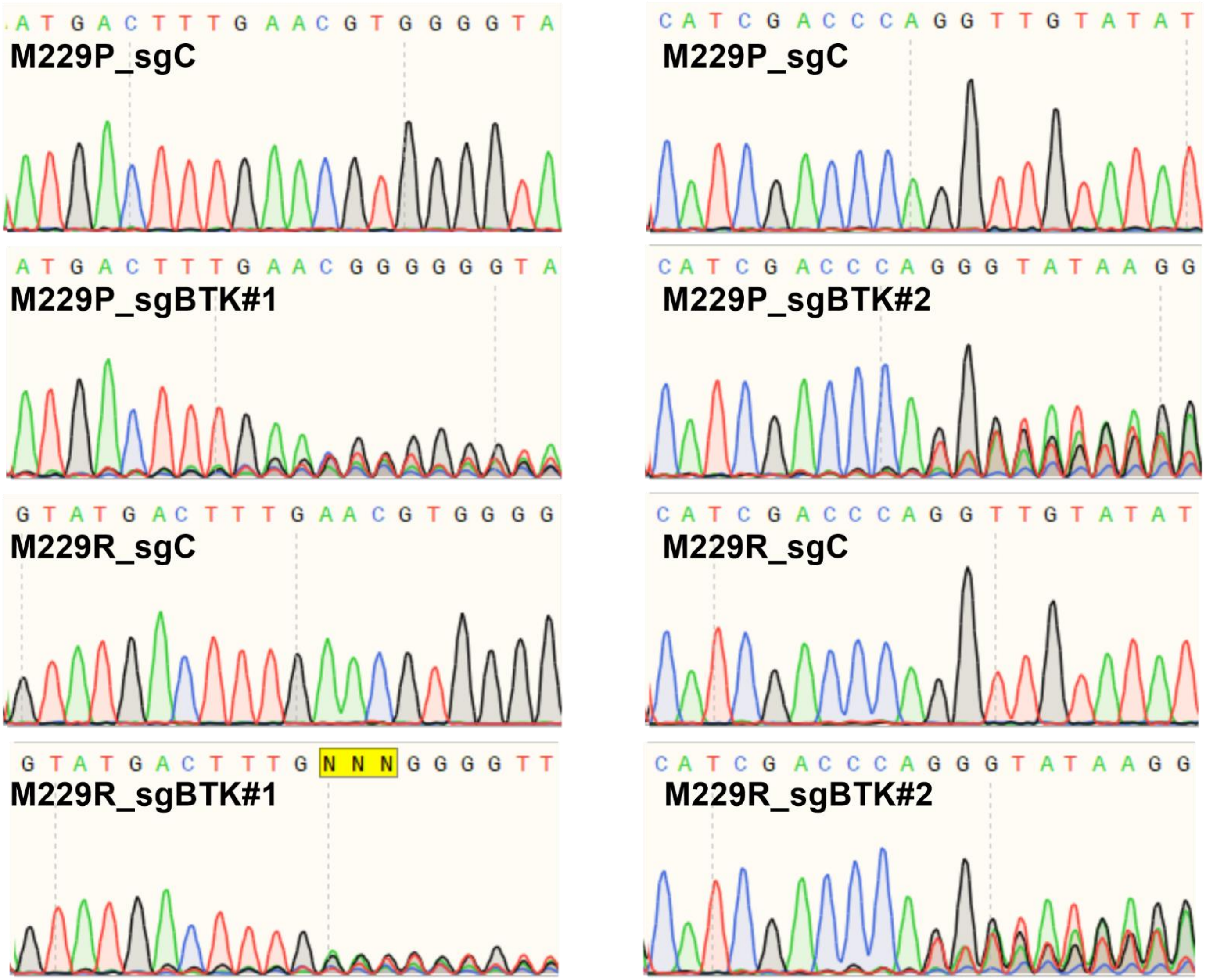


**Fig S9. Quantification of BTK knockout efficiency.** Representative Sanger sequencing traces that were used to measure CRISPR knockout efficiency with the TIDE algorithm as described in the Materials and Methods section.

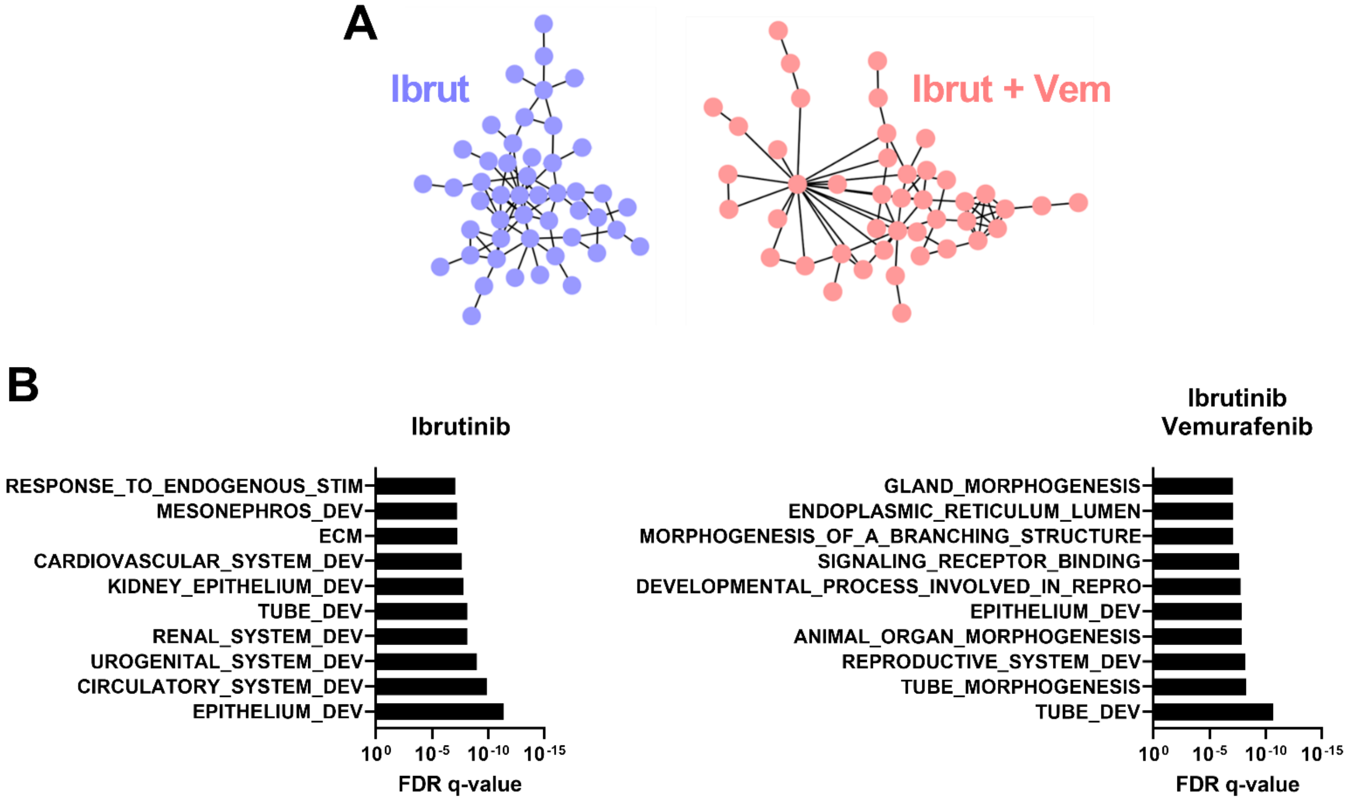


**Fig S10. Differential gene expression networks are associated with developmental gene signatures. A.** All differentially expressed genes in the ibrutinib-treated group (n = 101) were analyzed by string network analysis (left, blue). A similar analysis was performed for the top 101 differentially expressed genes in the ibrutinib + vemurafenib combination treatment group (right, red). **B.** Gene ontology analysis of genes within the interaction networks from Fig. 3B.

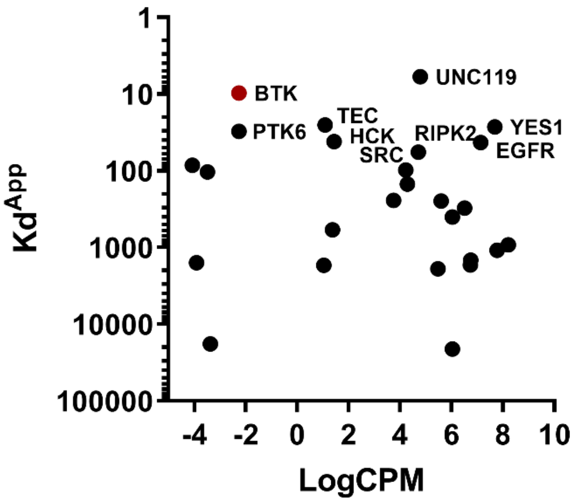


**Fig S11. Expression of ibrutinib targets in M229P/R cells.** Ibrutinib $K_d$ against various kinases (1) compared with kinase gene expression in M229P/R cells. RNA-seq data processing was performed as described in Materials and Methods.

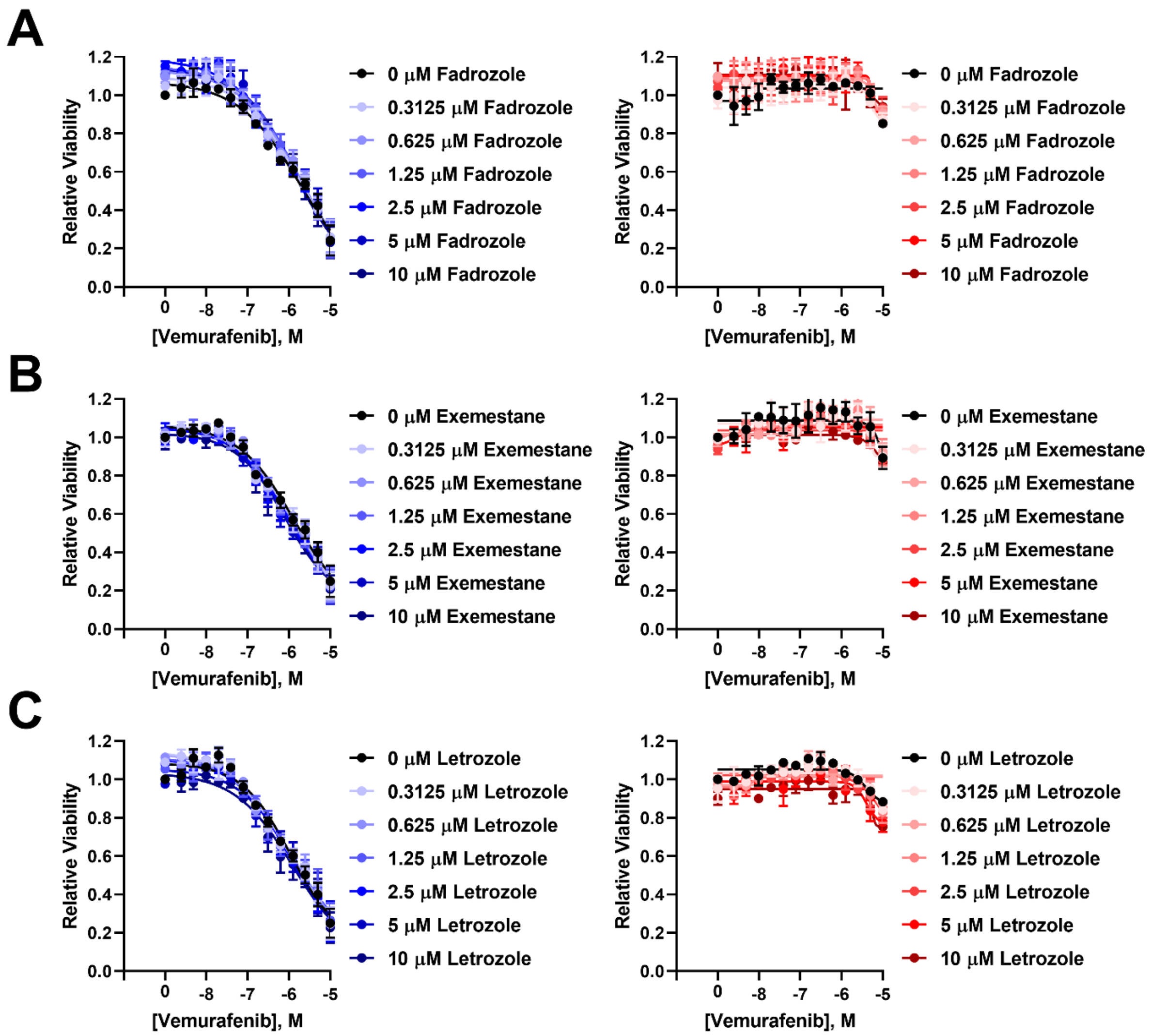


**Fig S12. Aromatase inhibitors do not alter BRAFi sensitivity.** M229P (blue) and M229R (red) cells were seeded into 384-well plates at a density of 1,000 cells/well. The next day the cells were treated with either **A.** Fadrozole, **B.** Exemestane, or **C.** Letrozole and vemurafenib as indicated. After 72 h viability was measured as described in Materials and Methods.

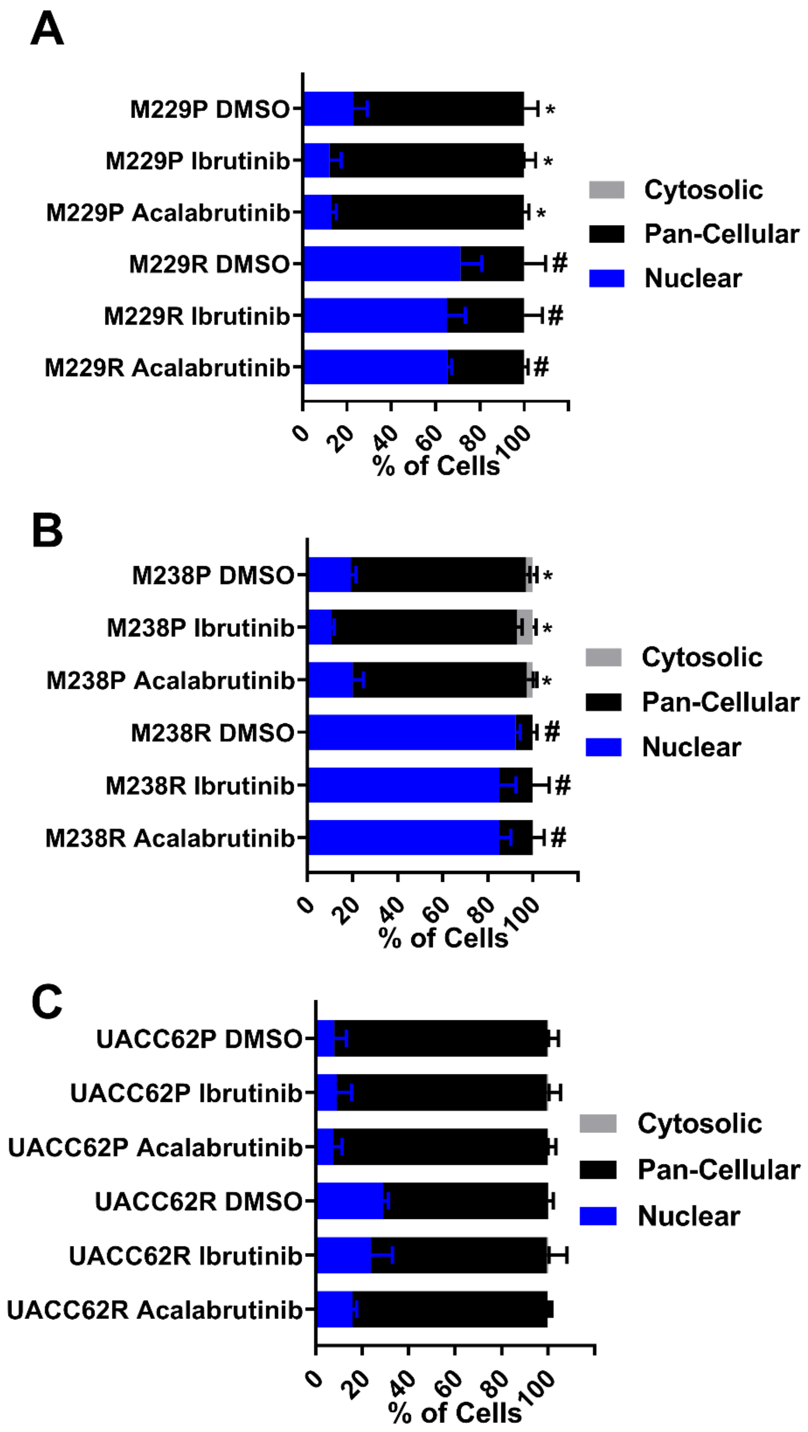


**Fig S13. Ibrutinib does not alter TAZ localization in BRAFi-resistant cells. A.** M229P/R, **B.** M238P/R, and **C.** UACC62P/R cells were seeded into 8-well chamber slides as described in Materials and Methods. The cells were treated with either DMSO, 5 µM ibrutinib, or 5 µM acalabrutinib. After 24 h the cells were fixed and stained as described in Materials and Methods. The proportion of cells with nuclear, pan-cellular, or cytosolic TAZ localization was quantified as described in Materials and Methods. Statistical analysis was performed on % of cells with nuclear localization where $p < 0.01$ was considered statistically significant. Bars marked with # indicate a statistically significant difference when compared with DMSO-treated parental cells and bars marked with * indicate a statistically significant difference when compared with DMSO-treated resistant cells.

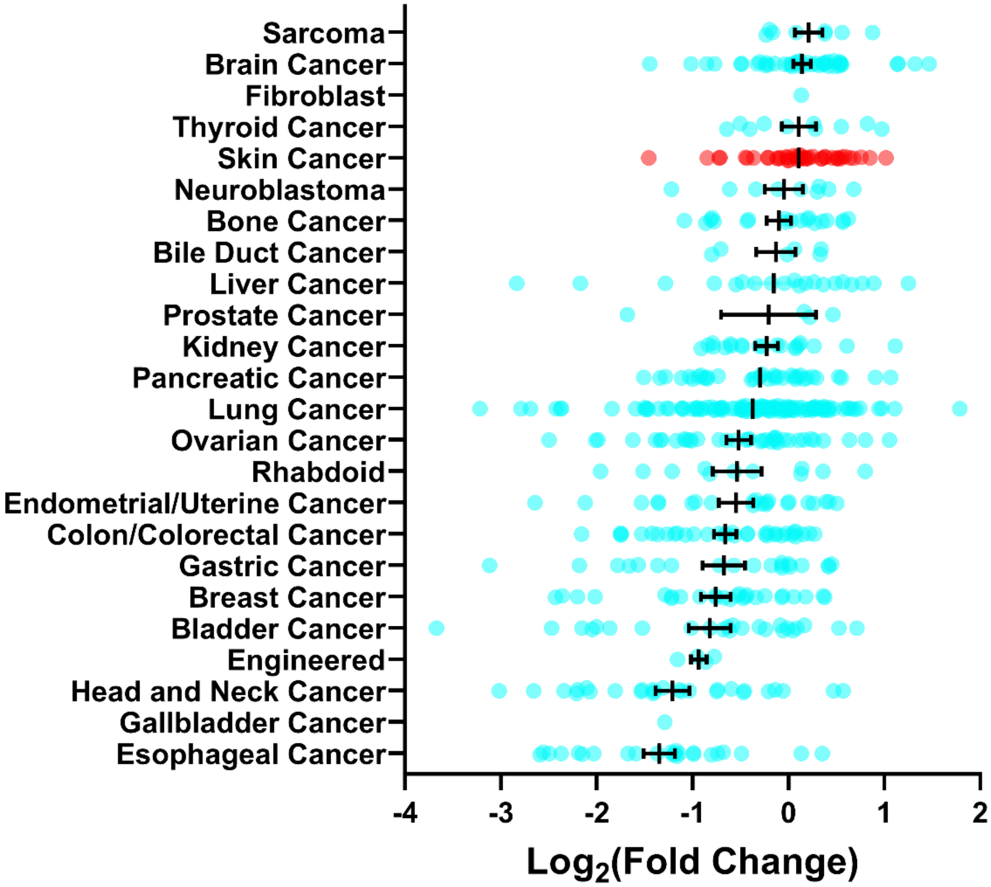


**Fig S14. Skin cancer cell lines are not sensitive to single agent ibrutinib treatment.** The cell lines in the PRISM dataset were stratified based on cancer type and ibrutinib sensitivity was compared. Smaller $Log_2$(Fold Change) values indicate higher sensitivity to ibrutinib.

1. Klaeger S, Heinzlmeir S, Wilhelm M, Polzer H, Vick B, Koenig PA*, et al.* The target landscape of clinical kinase drugs. Science **2017**;358
